# Structure and inhibition of the SARS-CoV-2 main protease reveals strategy for developing dual inhibitors against M^pro^ and cathepsin L

**DOI:** 10.1101/2020.07.27.223727

**Authors:** Michael Dominic Sacco, Chunlong Ma, Panagiotis Lagarias, Ang Gao, Julia Alma Townsend, Xiangzhi Meng, Peter Dube, Xiujun Zhang, Yanmei Hu, Naoya Kitamura, Brett Hurst, Bart Tarbet, Michael Thomas Marty, Antonios Kolocouris, Yan Xiang, Yu Chen, Jun Wang

## Abstract

The main protease (M^pro^) of SARS-CoV-2, the pathogen responsible for the COVID-19 pandemic, is a key antiviral drug target. While most SARS-CoV-2 M^pro^ inhibitors have a γ-lactam glutamine surrogate at the P1 position, we recently discovered several M^pro^ inhibitors have hydrophobic moieties at the P1 site, including calpain inhibitors II/XII, which are also active against human cathepsin L, a host-protease that is important for viral entry. To determine the binding mode of these calpain inhibitors and establish a structure-activity relationship, we solved X-ray crystal structures of M^pro^ in complex with calpain inhibitors II and XII, and three analogues of **GC-376**, one of the most potent M^pro^ inhibitors *in vitro*. The structure of M^pro^ with calpain inhibitor II confirmed the S1 pocket of M^pro^ can accommodate a hydrophobic methionine side chain, challenging the idea that a hydrophilic residue is necessary at this position. Interestingly, the structure of calpain inhibitor XII revealed an unexpected, inverted binding pose where the P1’ pyridine inserts in the S1 pocket and the P1 norvaline is positioned in the S1’ pocket. The overall conformation is semi-helical, wrapping around the catalytic core, in contrast to the extended conformation of other peptidomimetic inhibitors. Additionally, the structures of three **GC-376** analogues **UAWJ246**, **UAWJ247**, and **UAWJ248** provide insight to the sidechain preference of the S1’, S2, S3 and S4 pockets, and the superior cell-based activity of the aldehyde warhead compared with the α-ketoamide. Taken together, the biochemical, computational, structural, and cellular data presented herein provide new directions for the development of M^pro^ inhibitors as SARS-CoV-2 antivirals.

## INTRODUCTION

The COVID-19 pandemic emerged in late December 2020 in Wuhan, China and evolved to be one of the worst public health crisis in modern history. The impact of COVID-19 on global public health and economy has been severe. The etiological agent of COVID-19 is SARS-CoV-2, which shares ~78% genetic similarity with SARS-CoV, the virus that led to the SARS outbreak in 2003. Although coronavirus outbreaks such as COVID-19 are not unpredicted, the high mortality rate and the ease of transmission of SARS-CoV-2 is unprecedented.

Currently there are few antivirals and no vaccines available for SARS-CoV-2. As such, it is imperative to identify drug targets that could lead to effective antivirals. Guided by research of the similar coronaviruses, SARS-CoV and MERS-CoV, several viral proteins have been prioritized as SARS-CoV-2 antiviral drug targets: the spike protein, the RNA-dependent RNA polymerase (RdRp), the main protease (M^pro^), and the papain-like protease (PL^pro^).^1,2^ The SARS-CoV-2 RdRp inhibitor remdesivir was granted emergency use authorization from FDA on May 1^st^ 2020. Remdesivir has broad-spectrum antiviral activity against SARS-CoV, SARS-CoV-2, and MERS-CoV in cell culture.^3–5^ The antiviral efficacy was further confirmed in MERS-CoV infection mouse and rhesus macaque models.^6,7^ Additional RdRp inhibitors under investigation for SARS-CoV-2 include EIDD-2801, favipiravir (T-705), ribavirin, and galidesivir.^8,9^ The fusion inhibitor EK1C4, which was designed based on the H2 peptide in the S2 domain of the HCoV-OC43 spike protein, showed promising broad-spectrum antiviral activity against SARS-CoV-2, SARS-CoV, MERS-CoV, as well as human coronaviruses HCoV-229E, HCoV-NL63, and HCoV-OC43.^10,11^ Meanwhile, the M^pro^ has been extensively explored as a drug target for not only SARS-CoV-2, but also SARS-CoV, MERS-CoV, as well as enteroviruses, rhinoviruses, and noroviruses.^12^ M^pro^ is a viral encoded cysteine protease that has a unique preference for a glutamine residue at the P1 site in the substrate, which was recently confirmed for SARS-CoV-2 by substrate profiling.^13^ Consequently, the majority of designed M^pro^ inhibitors contain either 2-pyrrolidone or 2-piperidinone at the P1 site as a mimetic of the glutamine residue in the substrate.^14^ Examples include compounds **N3**, **13b**, **11a**, **11b**, and our recently identified **GC-376**,^15–18^ all of which have potent enzymatic inhibition in biochemical assay and antiviral activity in cell culture. Their mechanism of action and mode of inhibition were revealed by the drug-bound X-ray crystal structures.^15–18^

Interestingly, our previous study discovered two un-conventional SARS-CoV-2 M^pro^ inhibitors, calpain inhibitors II and XII, that are structurally dissimilar to the traditional M^pro^ inhibitors, such as **GC-376**.^15^ Specifically, calpain inhibitors II and XII incorporate the hydrophobic methionine and norvaline side chains in the P1 position. This discovery challenges the idea that a hydrophilic glutamine mimetic is required at the P1 position. Furthermore, calpain inhibitor II is a potent inhibitor of human protease cathepsin L, with a Ki of 50 nM.^19^ Cathepsin L plays an important role in SARS-CoV-2 viral entry by activating the viral spike protein,^20,21^ ^22^ and has a relatively broad substrate preference at the P1 position on the substrate.^23,24^ Studies have indicated that cathepsin L inhibitors can block or significantly decrease virus entry.^20,25^ To dissect the mechanism of action of these two promising drug candidates, we solved the high-resolution X-ray crystal structures of M^pro^ with calpain inhibitors II and XII (Fig. 1). We found that calpain inhibitor II is bound to M^pro^ in the canonical, extended conformation, but calpain inhibitor XII adopted an unexpected binding mode, where it assumes an inverted, semi-helical conformation in which the P1’ pyridine ring is placed in the S1 pocket instead of the P1 norvaline sidechain, as one would expect. The complex structures of calpain inhibitors II and XII, together with the structure-activity relationship studies of calpain inhibitors II/XII, revealed the S1 pocket of M^pro^ can accommodate both hydrophilic and hydrophobic substitutions, paving the way for the design of dual inhibitors that target both the viral M^pro^ and host cathepsin L. Finally, guided by the X-ray crystal structure of SARSA-CoV-2 M^pro^ with **GC-376** (PDB: 6WTT),^15^ three analogs **UAWJ246**, **UAWJ247**, and **UAWJ248** were designed to profile the sidechain preferences of the S1’, S2, S3 and S4 pockets. The X-ray crystal structures and activity profile presented herein offer valuable insights into the substrate promiscuity of M^pro^, as well as the design of new SARS-CoV-2 M^pro^ inhibitors.

**Fig. 1.**
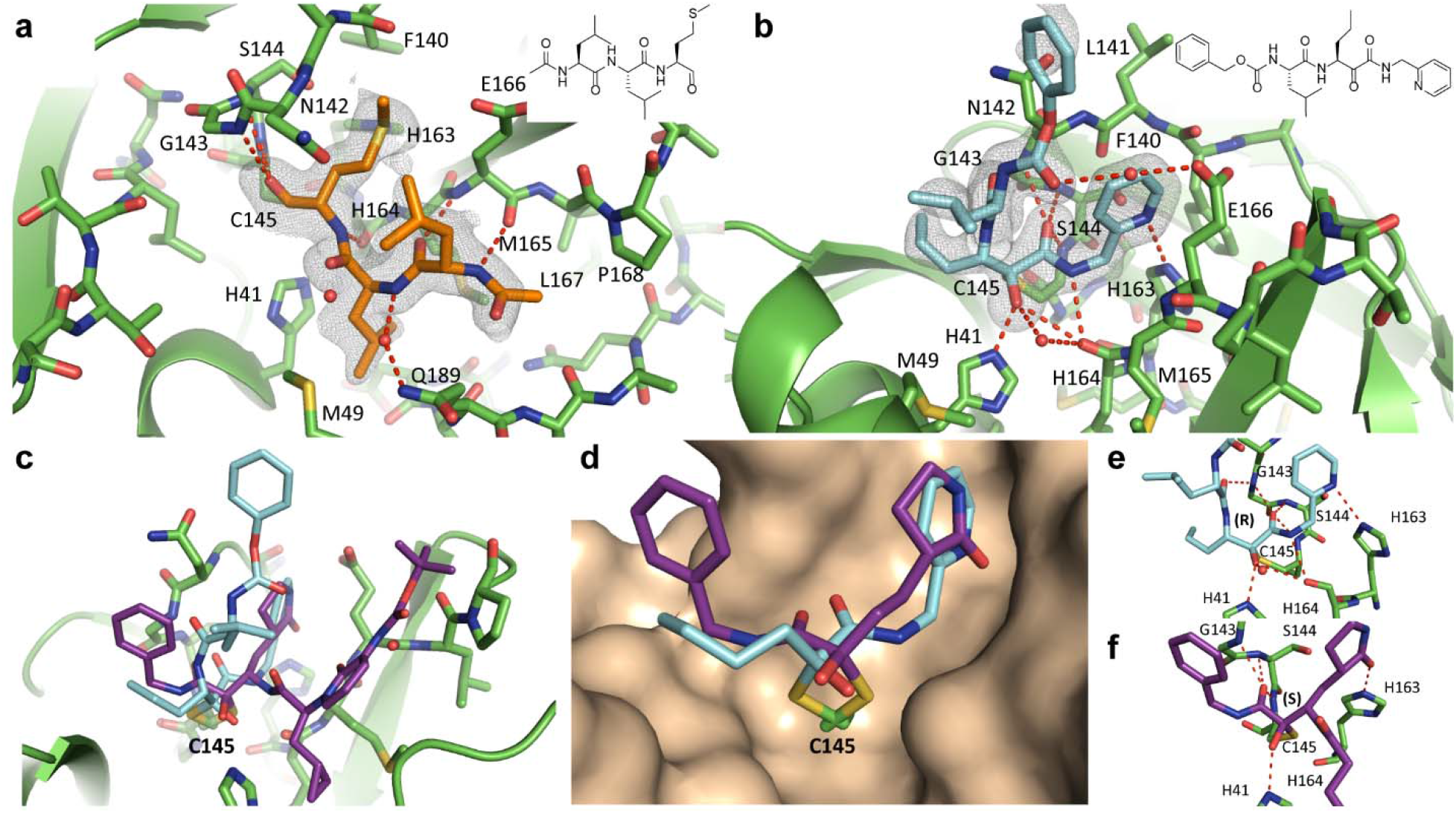
X-ray crystal structures of SARS-CoV-2 in complex with calpain inhibitors II (PDB: 6XA4) and XII (PDB: 6XFN). Hydrogen bonds are shown as red dashed lines. SARS-CoV-2 M^pro^ in complex with **(a)** calpain inhibitor II (orange) and **(b)** calpain inhibitor XII (blue). Unbiased Fo-Fc electron density map, shown in grey, is contoured at 2 σ. **c** Comparison of calpain inhibitor XII (blue) and SARS-CoV-2 M^pro^ α-ketoamide inhibitor **13b** (PDB ID 6Y2F) shown in purple. **d** Close-up comparison of calpain inhibitor XII (blue) and **13b** (purple) in the S1 and S1’ sites. The P1 pyrrolidinone ring and P1’ benzene of **13b** occupy the S1 and S1’ sites respectively. Conversely, the P1 norvaline and P1’ pyridine of calpain inhibitor XII adopt the S1’ and S1 sites. **e** Calpain inhibitor XII hydrogen bonding network in the catalytic core, accompanying a stereochemical inversion of the thiohemiketal adduct with Cys145, assuming the (R) configuration. **f** Hydrogen bonding network of **13b** in the catalytic core. Like all other α-ketoamide inhibitors, the covalent adduct adopts an (S) conformation.

## RESULTS AND DISCUSSION

### SARS-CoV-2 M^pro^ constructs used in this study

Three M^pro^ constructs were used in this study: the tag-free native M^pro^ (M^pro^), M^pro^ with two extra residues: histidine and methionine at the N-terminus (HM-M^pro^), the M^pro^ with a native N-terminus, and a C-terminal his-tag (M^pro^-His). The tag-free M^pro^ was used for all functional assays, while the other two constructs were used for structure determination due to ease of crystallization. The HM-M^pro^ construct was used for the complex structures of all five compounds including calpain inhibitors II and XII, **UAWJ246**, **UAWJ247**, and **UAWJ248**. In addition, the M^pro^-His construct was also co-crystallized with **UAWJ246** as a control for the potential influence of extra HM residues of the N-terminus.

The M^pro^-His and the native M^pro^ have similar enzymatic activity with *k*_cat_/K_m_ values of 6,689 s^−1^M^−1^, 5,748 s^−1^M^−1^, respectively (Supplementary Fig. S1a). The HM-M^pro^ construct has significantly reduced enzymatic activity with a *k*_cat_/K_m_ value of 214 s^−1^M^−1^, which is about 3.7% of the M^pro^ (*k*_cat_/K_m_ = 5,748 s^−1^M^−1^) (Supplementary Fig. S1a). This was expected as it has been shown that M^pro^ requires dimerization to be catalytically active and the N-terminal finger plays an essential role in dimerization.^26^ Specifically, the first residue serine (Ser1) from one protomer interacts with the Glu166 of the adjacent protomer, a feature that is important for catalytic activity (Supplementary Fig. S1b). Nevertheless, the HM-M^pro^ turned out to be an excellent construct for crystallization, and we were able to determine several high-resolution drug-bound X-ray crystal structures. In contrast, efforts to obtain high-quality crystals with the M^pro^-His construct was challenging, because of the localization of the disordered His-tag at the crystal packing interface.^15^ To validate the relevance of the use of the enzymatic inactive HM-M^pro^ construct for the crystallization, we crystalized **UAWJ246** with both the HM-M^pro^ and M^pro^-His constructs, and the binding pose of **UAWJ246** in these two constructs was nearly superimposable (Supplementary information, S3). Several previous studies similarly used the enzymatically inactive M^pro^ with extra residues at the N-terminus for the structural studies, and the ligand binding poses were identical to those with tag-free M^pro^ (e.g., PDB 7BRP vs 6WNP, 6WTJ vs 6L70).^27^ Therefore, the use of enzymatic inactive HM-M^pro^ construct for crystallographic study of inhibitor binding is justified.

### X-ray crystal structures of SARS-CoV-2 M^pro^ in complex with calpain inhibitors II and XII

Previous studies have shown SARS-CoV and SARS-CoV-2 M^pro^ cleave polyproteins at P2-P1 ↓ P1’ where P1’ is a residue with a small side chain (Ala, Ser, or Gly), P1 is glutamine and P2 is a large, hydrophobic residue, such as leucine or phenylalanine.^13,28,29^ This consensus sequence has operated as the foundation for extensive inhibitor designs where a reactive warhead, usually an aldehyde, α,β-unsaturated ester, or α-ketoamide, is linked to a glutamine surrogate pyrrolidone that is connected to a hydrophobic residue via an amide bond.^12,30–32^ This strategy has been largely successful, with the development of inhibitors such as **GC-376** and **13b** that have IC_50_ values in the low nM range for SARS-CoV-2 M^pro^.^15,17^

We recently reported calpain inhibitors II and XII and boceprevir have low μM IC_50_ values against SARS-CoV-2 M^pro^.^15^ Importantly, these compounds have a hydrophobic side chain at the P1 position, challenging the notion that a hydrophilic moiety is required at this position (Table 1). This has previously been demonstrated with SARS-CoV M^pro^, where the aldehyde inhibitor Cm-FF-H binds with a Ki of 2.24 ± 0.58 μM despite having a phenylalanine at the P1 position.^33^ The observed substrate plasticity is in part attributed to the reactivity of the electrophilic warhead aldehyde with the catalytic cysteine, which offsets the requirement for favorable interactions with the hydrophilic S1 subsite. Furthermore, it introduces the prospect of modifying the P1 residue so that it interacts with multiple host or viral protease enzymes that are essential for promoting SARS-CoV-2 viral entry or replication, which would increase antiviral spectrum and genetic barrier to resistance. To visualize the interactions between the P1 site and the hydrophobic S1 residue, we solved the complex structures of SARS-CoV-2 M^pro^ with calpain inhibitor II and calpain inhibitor XII.

**Table 1:**
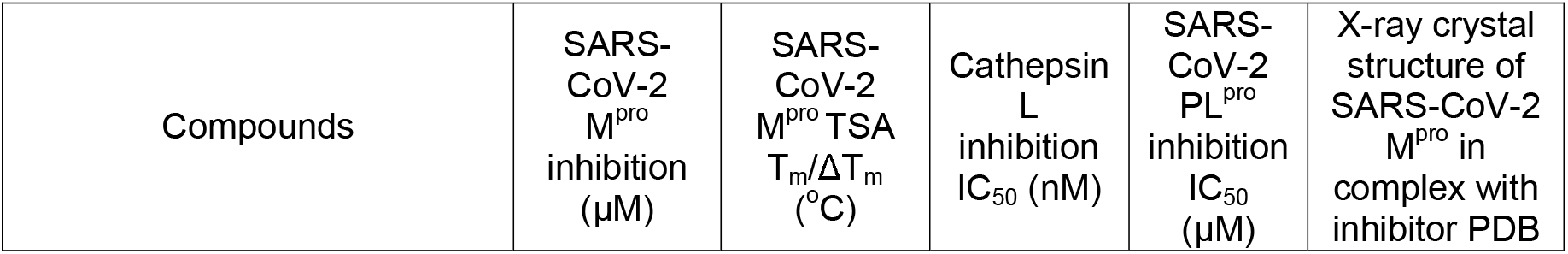
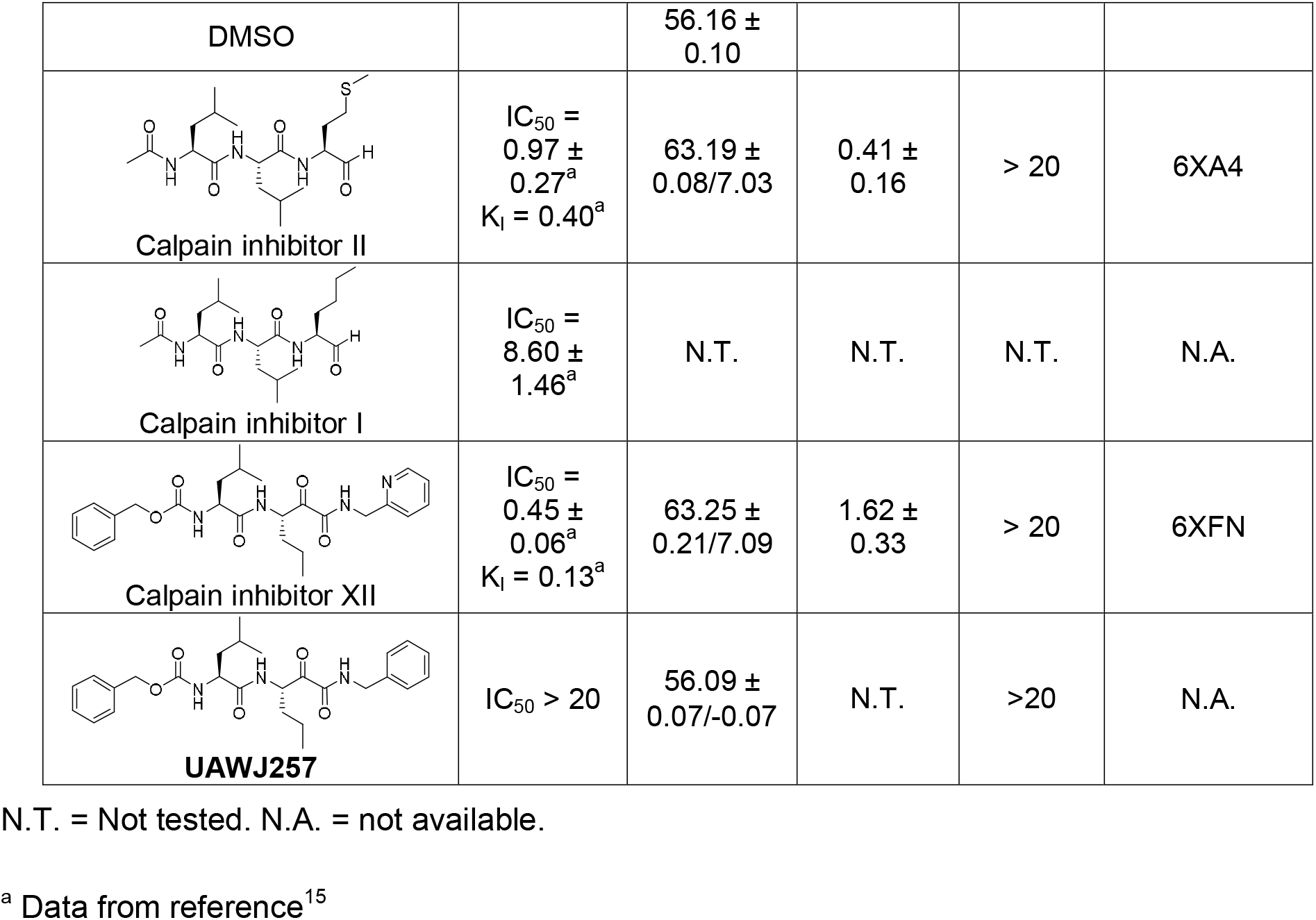
Biochemical and biophysical characterization of calpain inhibitors II and XII and their analogues as SARS-CoV-2 M^pro^ inhibitors.

| Compounds | SARS-CoV-2 M<sup>pro</sup> inhibition (μM) | SARS-CoV-2 M<sup>pro</sup> TSA T<sub>m</sub>/ΔT<sub>m</sub> (°C) | Cathepsin L inhibition IC<sub>50</sub> (nM) | SARS-CoV-2 PL<sup>pro</sup> inhibition IC<sub>50</sub> (μM) | X-ray crystal structure of SARS-CoV-2 M<sup>pro</sup> in complex with inhibitor PDB |
| --- | --- | --- | --- | --- | --- |
| DMSO |  | 56.16 ± 0.10 |  |  |  |
| Calpain inhibitor II | $IC_{50} = 0.97 \pm 0.27^a$<br>$K_i = 0.40^a$ | 63.19 ± 0.08/7.03 | 0.41 ± 0.16 | > 20 | 6XA4 |
| Calpain inhibitor I | $IC_{50} = 8.60 \pm 1.46^a$ | N.T. | N.T. | N.T. | N.A. |
| Calpain inhibitor XII | $IC_{50} = 0.45 \pm 0.06^a$<br>$K_i = 0.13^a$ | 63.25 ± 0.21/7.09 | 1.62 ± 0.33 | > 20 | 6XFN |
| **UAWJ257** | $IC_{50} > 20$ | 56.09 ± 0.07/-0.07 | N.T. | >20 | N.A. |

N.T. = Not tested. N.A. = not available.
<sup>a</sup> Data from reference<sup>15</sup>

The crystal structures of the SARS-CoV-2 HM-M^pro^ in complex with calpain inhibitors II (PDB: 6XA4) and XII (PDB: 6XFN) were solved in the C2 space group at 1.65 and 1.70 Å resolution respectively, with one protomer per asymmetric unit (Fig. 1, Supplementary Table 1). Like other peptidomimetic aldehyde inhibitors, the thiohemiacetal of calpain inhibitor II occupies the oxyanion hole formed by the backbone amide groups of Gly143, Ser144, and Cys145 (Fig. 1a). Here it adopts the (S) configuration, which is typical for most M^pro^ aldehyde inhibitors, although the (R) configuration has also been observed.^15,34^ Like previous M^pro^ and cathepsin L complex structures, the body of the inhibitor extends the length of the substrate-binding channel, with the side chains placed in their respective recognition pockets. The P1 methionine side chain projects into the S1 subsite where the sulfur forms a hydrogen bond (HB) with His163. The P2 valine side chain forms hydrophobic interactions in the S2 pocket, while the P3 valine occupies the solvent-accessible S3 position. Multiple hydrogen bonds form between the amide backbone and the main chains of His164, Met165, and Glu166.

In contrast to the pose of calpain inhibitor II, calpain inhibitor XII demonstrates an atypical binding mode where it adopts an inverted, semi-helical conformation that wraps around the catalytic core (PDB: 6XFN) (Fig. 1b). This is dissimilar to the extended configuration of previously published peptidomimetic inhibitors, including other α-ketoamide compounds such as **13b** (Fig. 1c, d).^18^ For calpain inhibitor XII, the P1’ pyridine is placed in the S1 site while the P1 norvaline occupies the S1’ site, the P2 valine projects outwards towards the solvent near the TSEDMLN loop (residues 45-51) and the terminal carboxybenzyl group curls back towards the S1 site, forcing Asn142 upwards while forming a water-mediated hydrogen bond with Glu166. Corresponding to this unique binding pose, we observed the (R) configuration of the thiohemiketal-Cys145 adduct for the first time among the known α-ketoamide inhibitors (Fig. 1d, e). From our recent X-ray crystal structure of the SARS-CoV-2 M^pro^ in complex with **GC-376** (PDB: 6WTT),^15^ it is known that the thiohemiacetal center of aldehyde-based inhibitors can assume either an (R) or (S) configuration, depending on which face of the aldehyde group undergoes nucleophilic attack from the thiolate of Cys145 during covalent bond formation.^15^ In contrast, the thiohemiketal group of all crystallographic solved α-ketoamide M^pro^ inhibitors such as **N3** and **13b** adopt the (S) configuration (Fig. 1f).^16,17^ The new calpain inhibitor XII structure demonstrates that, like aldehyde-based inhibitors, the covalent adduct formed between α-ketoamide compounds and the catalytic cysteine can assume two different configurations as well.

In both the (R) and (S) configurations of the thiohemiketal adducts, the hydroxyl group is placed near His41, while the amide oxygen is positioned in the oxyanion hole. However, the exact locations of these two functional groups result in different HB patterns. Compared to other α-ketoamide inhibitors, the unique binding mode of calpain inhibitor XII alters the hydrogen-bonding network of the catalytic core (Fig. 1e, f). The hydroxyl group forms a short HB (2.5 Å in length) with the catalytic His41, and two weak HBs (3.3 Å) with the main chain carbonyl of His164 and a water molecule in the central channel between the S1 and S2 pockets. The ketoamide amide oxygen establishes three HBs in the oxyanion hole (2.9, 3.2 and 3.1 Å to the backbone –NH of Gly143, Ser144 and Cys145 respectively), and its nitrogen forms a HB (3.1 Å) with the mainchain carbonyl of His164. In the canonical binding conformation for α-ketoamide inhibitors, such as **13b** and those described herein, the hydroxyl group establishes one standard HB (2.8 Å) with His41 and another HB (3.3 Å) with a bulk water molecule. Meanwhile, the HBs between the amide oxygen and the oxyanion hole now have distances of 3.3, 3.0 and 2.5 Å respectively, while the amide nitrogen forms no HBs with the protein.

The S1 pocket recognizes the most conserved residue in the M^pro^ substrate, the P1 glutamine. Underscoring its importance for ligand binding, most specific M^pro^ inhibitors have a glutamine surrogate such as the pyrrolidone ring that occupies the S1 pocket. Compounds like calpain inhibitor II, calpain inhibitor XII, and Boceprivir prove that a hydrophobic side chain can also be accommodated in the S1 pocket. Like previous inhibitors, hydrophobic interactions are observed between the newly identified inhibitors and the backbone atoms of Leu141/Asn142/Met165. However, these interactions are further enhanced in calpain inhibitor XII, where its aromatic pyridine ring is stacked and sandwiched between the two planar peptide bonds involving Asn142 and Met165 respectively. Furthermore, a hydrogen bond is observed between His163 and the methionine sulfur and pyridine nitrogen for calpain inhibitors II and XII, respectively.

To dissect the importance of the hydrogen bond between His163 and the pyridine nitrogen from calpain inhibitor XII, we designed the benzene counterpart of calpain inhibitor XII, compound **UAWJ257**. It was found that **UAWJ257** demonstrated no detectable inhibition against SARS-CoV-2 M^pro^ (IC_50_ > 20 μM) (Table 1). Likewise, no binding was detected in the thermal shift assay (ΔT_m_ = −0.07 °C). In addition to abolishing the hydrogen bond, the dramatic loss of inhibition might be attributed to a clash between His163 and the benzene hydrogen that replaces the lone pair on the pyridine nitrogen, which lies only 3.1 Å away from His163. Similarly, this hydrogen bond is important for calpain inhibitor II binding, since its butyl analogue, calpain inhibitor I, has an IC_50_ that is ~ 10-fold weaker (Table 1). Overall, the structure-activity relationship results of calpain inhibitors II and XII are consistent with the binding poses shown in the X-ray crystal structures.

While calpain inhibitor II was previously reported to inhibit cathepsin L with an inhibition constant Ki of 0.6 nM,^19^ which was confirmed by our data as well (IC_50_ = 0.41 nM), the inhibition of cathepsin L by calpain inhibitor XII was unknown. We determined that calpain inhibitor XII is also a potent inhibitor of cathepsin L, with an IC_50_ value of 1.62 ± 0.33 nM. Because cathepsin L has been shown to activate the SARS-CoV-2 spike protein, cathepsin L inhibitors decrease viral entry.^20^ Importantly, this may provide an explanation for the superior antiviral activity of calpain inhibitor II and XII despite having inferior affinity for M^pro^ compared to the specific inhibitors **GC-376**, **N3**, **UAWJ246**, **UAWJ247**, and **UAWJ248** (Table 1, 2 and references^15,18^). Both calpain inhibitors II and XII had no detectable inhibition against the SARS-CoV-2 papain-like protease (PL^pro^) (IC_50_ > 20 μM) (Table 1), suggesting they are not non-specific cysteine protease inhibitors. Collectively, the X-ray crystal structures of SARS-CoV-2 HM-M^pro^ in complex with calpain inhibitors II and XII, along with the enzymatic assay results, suggest that it is feasible to develop dual inhibitors that simultaneously targeting the SARS-CoV-2 M^pro^ and the host cathepsin L, both of which are validated antiviral drug targets for SARS-CoV-2.^20,21^

**Table 2:**
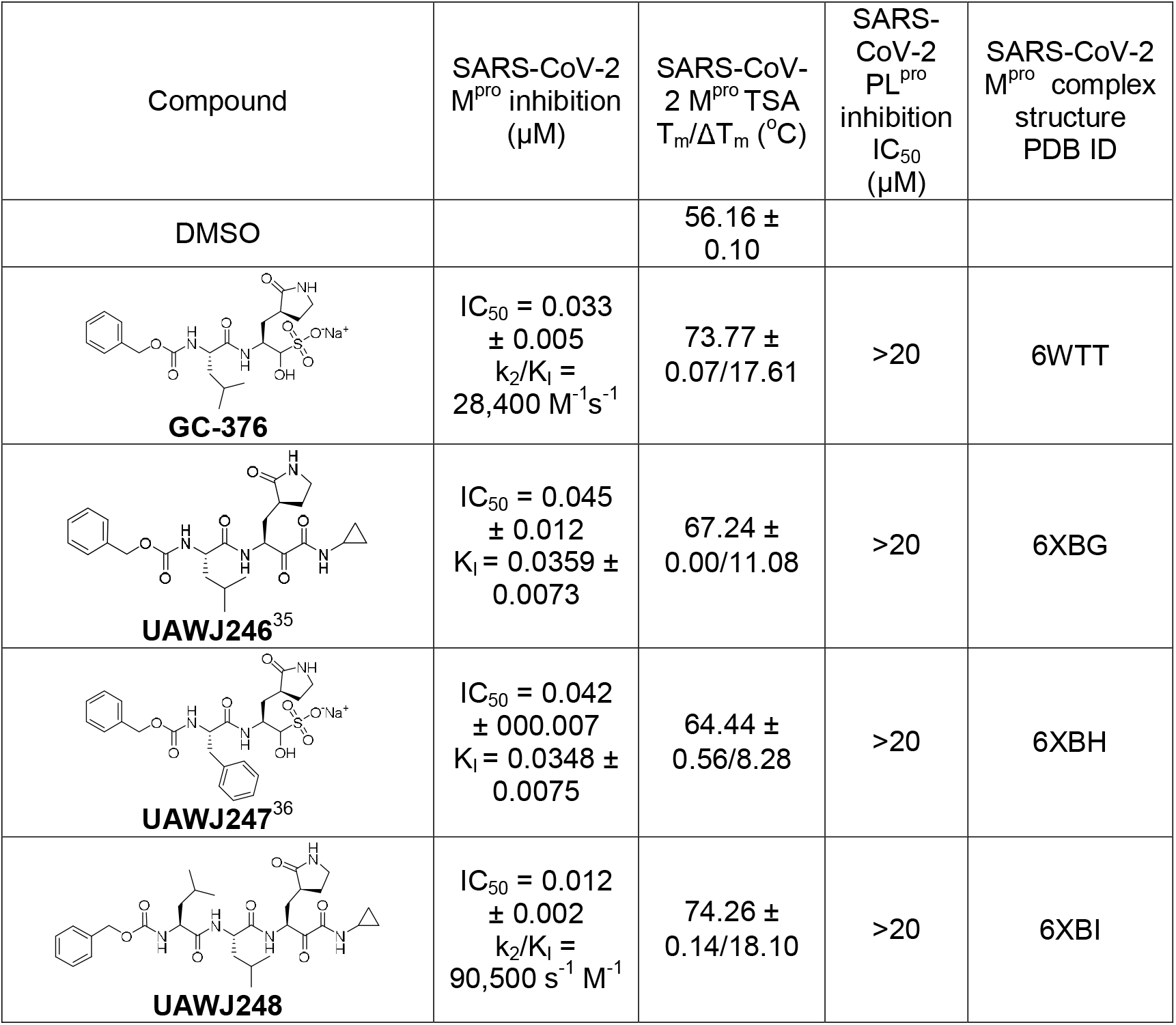
Biochemical and biophysical characterization of GC-376 analogues as SARS-CoV-2 M^pro^ inhibitors.

### Rational design of GC-376 analogues and the X-ray crystal structures of SARS-CoV-2 M^pro^ in complex with UAWJ246, UAWJ247, and UAWJ248

We recently demonstrated the inhibition of SARs-CoV-2 M^pro^ by **GC-376**, and solved the X-ray crystal structure of M^pro^-His with **GC-376** (PDB: 6WTT).^15^ To profile the substrate spectrum of SARS-CoV-2 M^pro^ in the S1’, S2, S3, and S4 sites, several **GC-376** analogues were designed. Specifically, compound **UAWJ246** was designed to occupy the S1’ pocket according to the overlay structures of SARS-CoV-2 M^pro^+**GC-376** (PDB: 6WTT) and SARS-CoV M^pro^ (H41A)+substrate (PDB: 2Q6G) (Supplementary Fig. S2). We found that **UAWJ246** inhibited SARS-CoV-2 M^pro^ with an IC_50_ value of 0.045 ± 0.012 μM, which is similar to that of GC-376 (IC_50_ = 0.033 ± 0.005 μM) (Table 2). **UAWJ246** contains a pharmacological compliant α-ketoamide reactive warhead with a cyclopropyl substitution. **UAWJ247** was designed to probe the substrate promiscuity in the S2 pocket. In the X-ray crystal structure of M^pro^-His with **GC-376**, the TSEDMLN loop constituting the S2 pocket exhibits significant flexibility among the three protomers in the asymmetric unit, indicating a variety of substitutions can be accommodated at this position. To test this hypothesis, **UAWJ247** was designed with a benzyl substitution at the P2 position instead of the isopropyl of **GC-376**. We found that **UAWJ247** has a similar inhibition constant of SARS-CoV-2 M^pro^ as **GC-376** (IC_50_ = 0.042 μM, Table 2). **UAWJ248** is a tripeptide and was designed to incorporate the P3 substitution. This slightly improved the IC_50_ to 0.012 μM, a ~3 -fold improvement compared to **GC-376** (Table 2). None of these compounds showed inhibition against the SARS-CoV-2 PL^pro^ (IC_50_ > 20 μM) (Table 2).

Next, we determined the mechanism of action with thermal shift binding assay, native mass spectrometry, and enzyme kinetic studies. As expected, binding of **UAWJ246**, **UAWJ247**, and **UAWJ248** all stabilized SARS-CoV-2 M^pro^, as shown by the ΔT_m_ shift of 11.08, 8.28, and 18.10 °C, respectively (Fig. 2a). All three compounds also stabilized the M^pro^-His construct to the same degree as the tag free M^pro^, suggesting these two constructs are functional equivalent (Fig. 2a). In contrast, compounds **UAWJ246** and **UAWJ247** did not show stabilization for the HM-M^pro^ construct, while the more potent **GC-376** and **UAWJ248** stabilized this construct with ΔT_m_ shift of 2.45, and 10.01 °C, respectively (Fig. 2a).

**Fig. 2.**
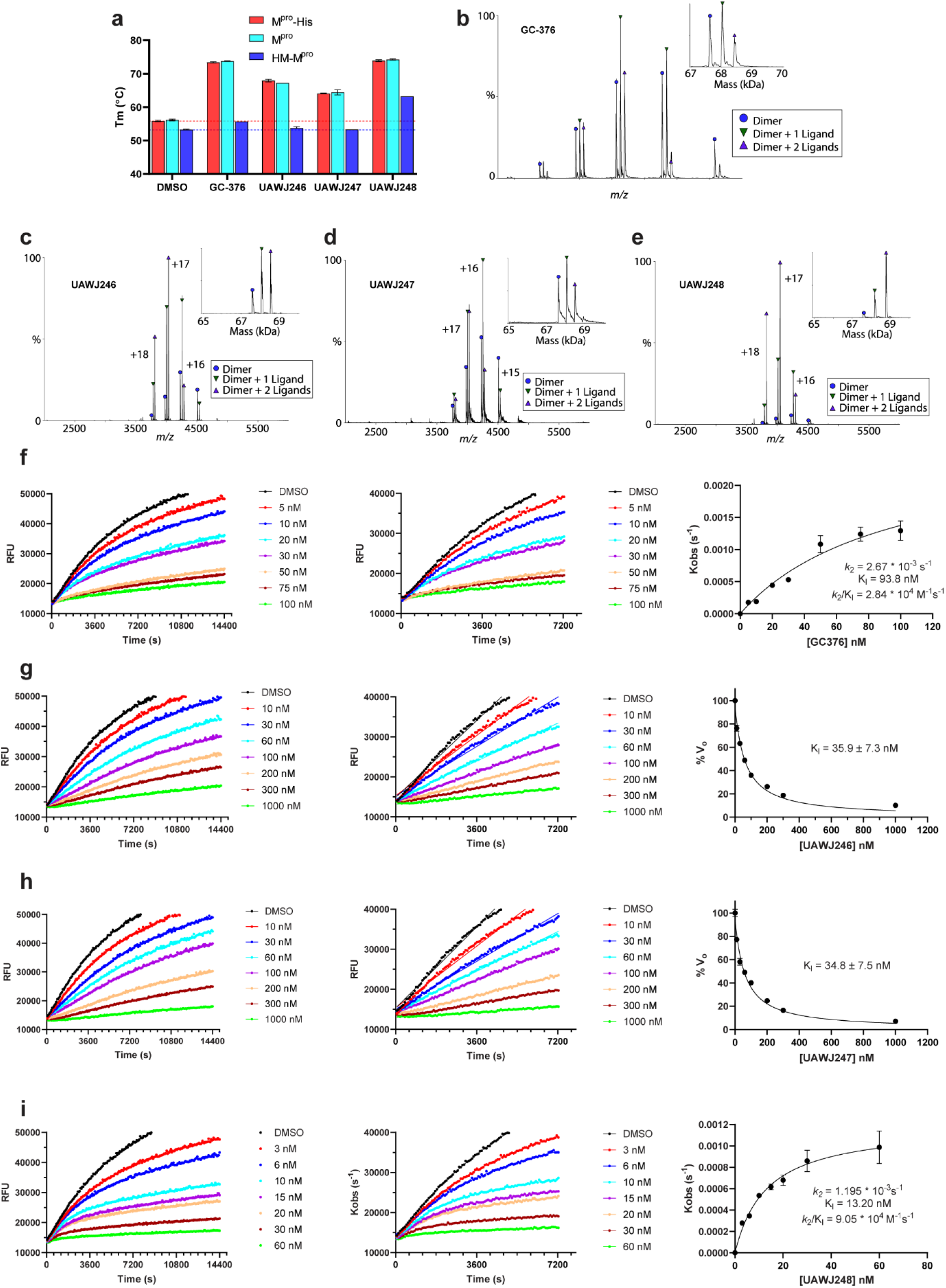
Pharmacological characterization of the mechanism of action of GC-376 analogues UAWJ246, UAWJ 247, and UAWJ 248 in inhibiting SARS-CoV-2 M^pro^. **a** Thermal shift binding assay of **GC-376** analogues with different SARS-CoV-2 M^pro^ constructs. Test compounds were incubated with 3 μM M^pro^ protein in reaction buffer (20 mM HEPES, pH 6.5, 120 mM NaCl, 0.4 mM EDTA, 4 mM DTT and 20% glycerol) at 30 °C for 30 min. 1X SYPRO orange dye was added and fluorescence of the well was monitored under a temperature gradient range from 20 °C to 90 °C with 0.05 °C/s incremental step. **b–e** Binding of inhibitors to SARS-CoV-2 M^pro^ using native mass spectrometry. Native mass spectra with the inset deconvolved spectra revealing ligand binding to with (**b**) 10 μM **GC-376** added, (**c**) 10 μM **UAWJ246**, (**d**) 10 μM **UAWJ247** added, and (**e**) 10 μM **UAWJ248** with 4 mM DTT added. The peaks are annotated with the blue circle as the dimer, green down triangle as the dimer with one ligand bound, and the purple up triangle as the dimer with two ligands bound. **f–i** Proteolytic reaction progression curves of SARS-CoV-2 M^pro^ in the presence or the absence of compounds. In the kinetic studies, 5 nM M^pro^ was added to a solution containing various concentrations of protease inhibitors and 20 μM FRET substrate to initiate the reaction. The reaction was then monitored for 4 h. The left column shows the reaction progression up to 4 h; middle column shows the progression curves for the first 2 h, which were used for curve fitting to generate the plot shown in the right column. Detailed methods were described in “Materials and methods” section. **GC-376** (**f**); **UAWJ246** (**g**); **UAWJ247** (**h**); **UAWJ248** (**i**).

The binding of all three compounds **UAWJ246**, **247**, and **248** to M^pro^ was further confirmed by native mass spectrometry (Fig. 2b-e). Like **GC-376**, addition of the ligands resulted in two new sets of peaks corresponding to one ligand per dimer and two ligands per dimer.

Enzyme kinetic studies showed that compounds **UAWJ246** and **UAWJ247** bound to M^pro^ reversibly with inhibition constant K_I_ values of 0.036 ± 0.007 and 0.035 ± 0.008 μM, respectively (Fig. 2g, h). In contrast, the enzyme kinetic curves for compound **UAWJ248** (Fig. 2i) was similar to that of **GC-376** (Fig. 2f), which showed a biphasic progression character, suggesting **UAWJ248** inhibits M^pro^ through a two-step process with an initial reversible binding followed by an irreversible inactivation. Fitting the progression curves with the two-step Morrison equation revealed the first step equilibrium dissociation constant K_I_ and the second step reaction constant *k_2_* as 13.20 nM and 0.001195 s^−1^, respectively, which corresponds to an overall *k_2_*/K_I_ value of 9.05 × 10^4^ M^−1^s^−1^. In comparison, the *k_2_*/K_I_ value for **GC-376** is 2.84 × 10^4^ M^−1^s^−1^, suggesting **UAWJ248is** 3.2-fold more potent than **GC-376**.

Most M^pro^ inhibitors with antiviral activity against SARS-CoV-2 use an α-ketoamide or aldehyde/bisulfite warhead to form a covalent adduct with the catalytic Cys145.^15,17^ We previously reported **GC-376** as one of the most potent inhibitors of SARS-CoV-2 M^pro^ in vitro with an IC_50_ of 0.033 μM and EC_50_ of 3.37 μM in the enzymatic assay and antiviral CPE assay.^15^ Here we show the α-ketoamide analogue of **GC-376**, **UAWJ246** has a comparable IC_50_ of 0.045 μM, suggesting the α-ketoamide and the aldehyde are nearly equivalent in terms of inhibitory activity. We solved the complex structure of **UAWJ246** with both SARS-CoV-2 HM-M^pro^ at 1.45 Å resolution as a dimer (PDB: 6XBG) and M^pro^-His at 2.35 Å as a trimer in the asymmetric unit (Fig. 3a, b; Supplementary information, Fig. S3). The binding pose of **UAWJ246** in these two constructs was nearly superimposable (Supplementary information, Fig. S3), suggesting the use of enzymatic inactive HM-M^pro^ construct for crystallographic study of inhibitor binding is justified. In the dimer structure of HM-M^pro^ with **UAWJ246**, the amide oxygen of the α-ketoamide occupies the oxyanion hole, while the thiohemiketal hydroxide forms a hydrogen bond with the catalytic histidine, His41. The cyclopropyl group extends partly into the S1’ subpocket. Interestingly, the carboxybenzyl of **UAWJ246** adopts a different conformation in each protomer (Fig. 3a, b). In the A protomer of the 1.45 Å resolution structure crystallized in P2_1_ space group, the carboxybenzyl moiety extends the length of the S3 and S4 subsites, forcing residues 189-191 to flip outwards (Fig. 3a). In the B protomer, the carboxybenzyl moiety projects upwards, where the S3 sidechain is normally positioned (Fig. 3b). The A-conformation resembles that of **GC-376** in our previously solved structure (PDB: 6WTT) (Supplementary information, Fig. S4a), whereas the B-conformation resembles **GC-376** from PDB ID 7BRR (Supplementary information, Fig. S4b) ^37^. The downward conformation establishes extensive interactions with the S4 pocket, and may be one of the main reasons for the superior *in vitro* activity of **GC-376** and its analogues. The upward conformation also forms favorable interactions with protein residues constituting the S3 site, including Glu166 and Gln189. Additionally, it enables the formation of intramolecular interactions between the benzyl ring and the P1 sidechain, similar to the hydrophobic intramolecular interactions formed between the P1 and P3 moieties in calpain inhibitor II and boceprevir, as well as, to some degree, the π-π stacking between the pyridine and the carboxybenzyl of calpain inhibitor XII (Fig. 2-3; Supplementary information, Fig. S5).^38^ It is likely that **GC-376**, **UAWJ246**, and other analogues exist in a dynamic equilibrium between these conformations and the captured crystallographic poses are, in part, determined by the crystal-packing interface between protomers and/or differences in the pH or ionic strength of the crystallization solution.

**Fig. 3.**
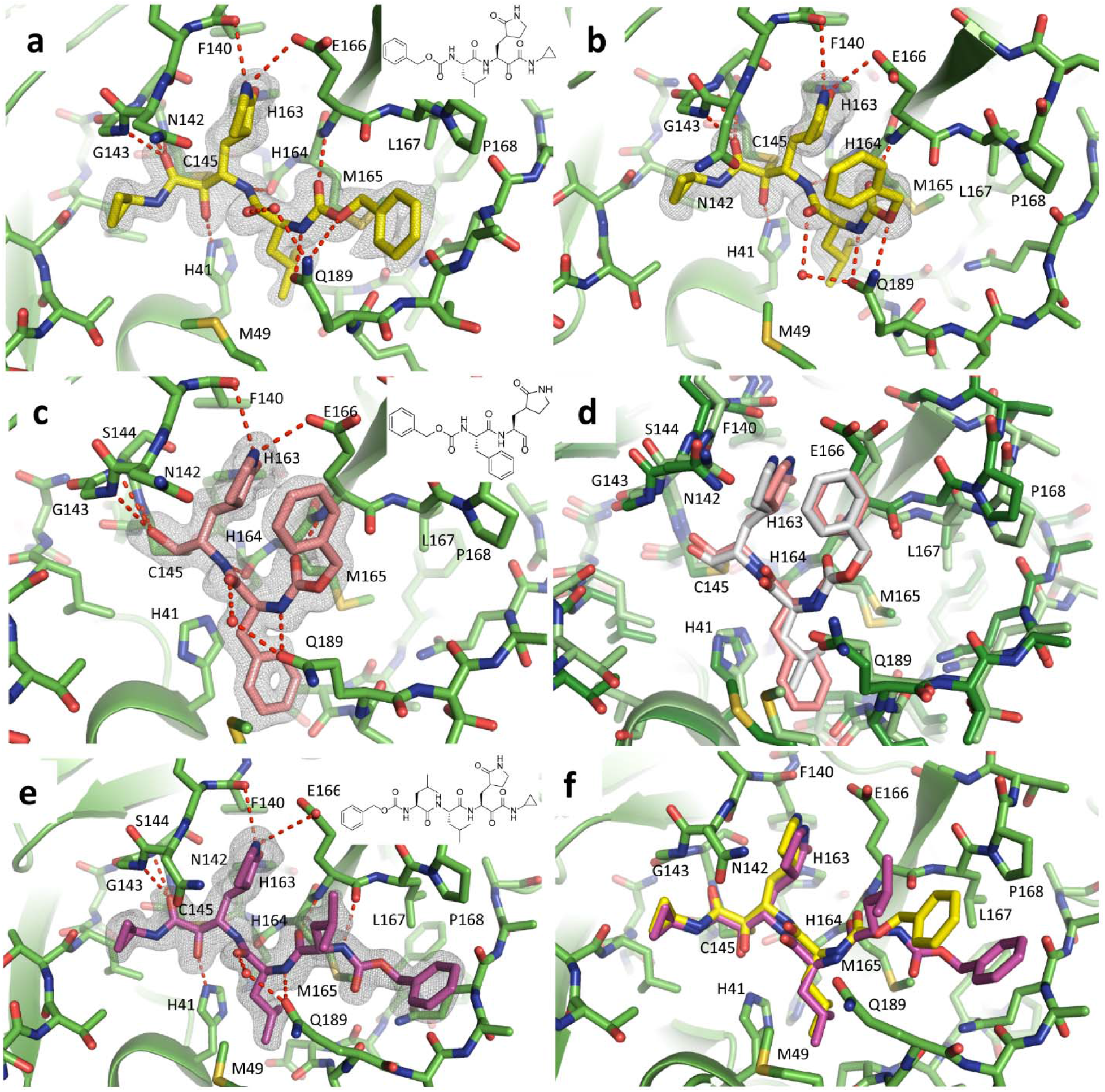
SARS-CoV-2 M^pro^ in complex with GC-376 analogues. Unbiased Fo-Fc electron density map, shown in grey, is contoured at 2 σ. Hydrogen bonds are shown as red dashed lines. Solved as a dimer in the P2_1_ spacegroup, we observe two different conformations of the carboxybenzyl group of **UAWJ246** in the **(a)** protomer A and **(b)** protomer B. **c** The complex structure of **UAWJ247**, revealing the P2 position can accommodate a Phe side chain. **D** Comparison of the binding poses of **UAWJ247** (dark green/salmon) and **GC-376** (light green/grey, PDB ID 7BRR). **e** The complex structure of **UAWJ248**, solved as a dimer in the P1 space-group. Protomer A is shown here, and the inhibitor binding pose is identical in protomer B. **f** Comparison of the binding poses of of **UAWJ248** (purple) and **UAWJ246** (yellow) in protomer A.

The chemical structure of **UAWJ247** is nearly identical to **GC-376**, except for the replacement of its S2 isobutyl moiety for a benzyl group, analogous to a Leu → Phe exchange. To visualize the binding mode of **UAWJ247**, we solved the complex structure with SARS-CoV-2 M^pro^ at 1.60 Å in the C2 space group with one protomer per asymmetric unit (PDB: 6XBH) (Fig. 3c, d). Like their chemical structures, the binding poses between **UAWJ247** and **GC-376** are very similar (Fig. 3d), with minor differences observed for Gln189 and the catalytic histidine, His41, which swivels towards the S2 benzyl group to form face-to-face π-stacking interactions. As expected, the IC_50_ of 0.045 μM for **UAWJ247** is very close to that of **GC-376** and consistent with the preference for a hydrophobic residue at the S2 site. This data also suggests replacing Leu for a larger Phe is tolerated, and that aromaticity can be incorporated into the S2 site for the purpose of improving pharmacokinetic properties or broadening the spectrum of activity, with limited effect on M^pro^ inhibition.

**UAWJ248** was designed to occupy the additional S4 pocket compared to **UAWJ246**. We solved the complex structure of **UAWJ248** with SARS-CoV-2 HM-M^pro^ at 1.70 Å as a dimer in the P1 monoclinic space group (PDB: 6XBI) (Fig. 3e, f). The conformation is consistent in both protomers The α-ketoamide warhead forms an adduct with Cys145 in the (S) conformation, like other cyclopropane α-ketoamide analogues described herein including **UAWJ246** and previously published **13b**.^17^ Similarly, the P1 γ-lactam and P2 isobutyl moieties occupy their respective S1 and S2 subsites. The P3 isobutyl orients upwards into the S3 site where it forms no meaningful interactions. However, the insertion of an additional leucine into the **UAWJ246** core ensures the formation of a hydrogen bond with the main chain amide oxygen of Glu166. The terminal carboxybenzyl is placed in the S4-S5 site, where nonspecific interactions occur between the benzene and side chain of Pro168 and Ala191 and π stacking with the main chain amides of Gln189, Thr190, and Ala191.

### Molecular dynamics simulations of SARS-CoV-2 M^pro^ with inhibitors

The binding interactions between the covalently bound calpain inhibitor II, calpain inhibitor XII, **UAWJ246**, **UAWJ247**, and **UAWJ248**, with SARS-CoV-2 M^pro^ were further explored using 100 ns-MD simulations with starting structures the X-ray structures with PBD IDs 6XA4, 6XFN, 6XBG (dimer), 6XBH (monomer), 6XBI (dimer). The simulations demonstrate that the complexes formed are stable, and the ligand positions do not deviate significantly from the crystallographic ones, with Cα RMSD values less than 2.4 Å and an overall ligand RMSD less than 3.5 Å (Fig. 4). The MD simulations further verified the stability of the interactions inside the binding cavity of SARS-CoV-2 M^pro^ observed in the X-ray structures, as inspected from the trajectories and shown in frequency interaction plots (Fig. 4a, d, g, j, m, p, s).

**Fig. 4.**
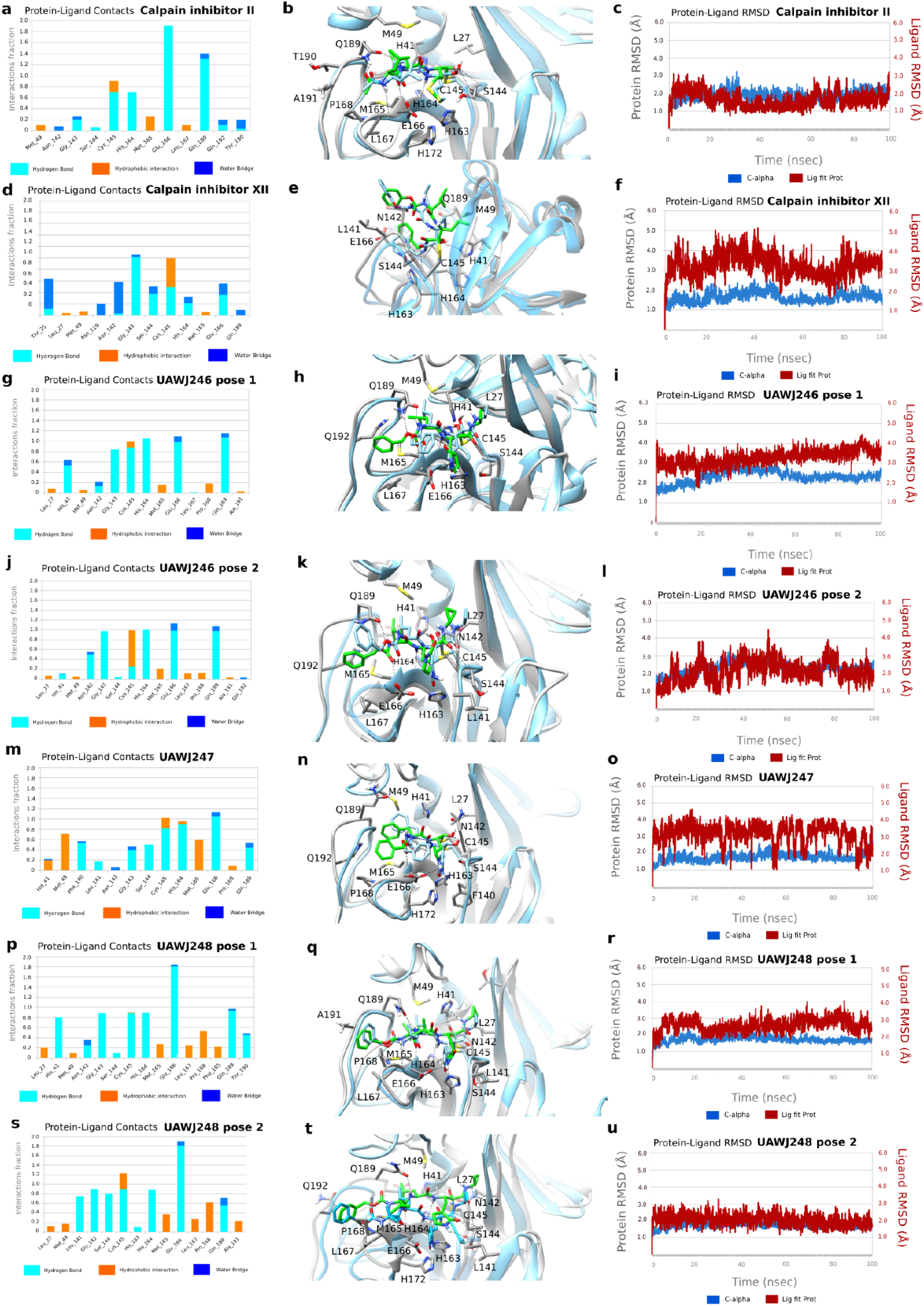
MD simulations of SARS-CoV-2 M^pro^ with its inhibitors. In (**a**), (**d**), (**g**), (**j**), (**m**), (**p**), (**s**) hydrogen bonding interactions bar is depicted in light blue, van der Waals in orange, water bridges in blue. Interactions are plotted from 100-ns MD simulations for the complexes between the covalently bound calpain inhibitor II, calpain inhibitor XII, **UAWJ246** (pose 1: first protomer), **UAWJ246** (pose 2: second protomer), **UAWJ247**, **UAWJ248** (pose 1: first protomer), **UAWJ248** (pose 2: second protomer) inside SARS-CoV-2 M^Pro^. They are considered important when frequency bar is ≥ 0.2. In (**b**), (**e**), (**h**), (**k**), (**n**), (**q**), (**t**) the last snapshots of the above mentioned 100ns-MD simulated complexes overlaid with experimental structures with PDB IDs 6XA4, 6XFN, 6XBG, 6XBH, 6XBI, respectively, are shown. In (**c**), (**f**), (**i**), (**l**), (**o**), (**r**), (**u**) the RMSD plots of Cα carbons (blue diagram, left axis) and of ligand (red diagram, right axis) of the above mentioned 100ns-MD simulated complexes are shown.

Compared to calpain inhibitor II (Fig. 4a, b), in **UAWJ247**, which also has an aldehyde warhead, the methionine P1 substituent was changed to 3-(2-pyrrolidinone) and additional hydrogen bonding interactions are formed. **UAWJ247** forms important hydrogen bonding interactions between the P1 2-pyrrolidinone NH group and E166 side chain, and peptidic carbonyl of F140 (Fig. 4m, n), in addition to the hydrogen bonds with G143, S144, C145, H164, Q189. Furthermore, the P2 benzyl group in **UAWJ247** fits better in the S2 subsite than the isopropyl from calpain inhibitor II, resulting in new van der Waals interactions with M49, H41, M165 (Fig. 4m, n). These additional stabilizing interactions reduce the IC_50_ against SARS-CoV-2 M^pro^ by 23-fold, i.e., from 0.97 μΜ in calpain inhibitor II to 0.042 μΜ in **UAWJ247**.

Calpain inhibitor XII, **UAWJ246** and **UAWJ248** are ketoamides having a ketone carbonyl compared to the aldehyde group in calpain inhibitor II and **UAWJ247**. Calpain inhibitor XII forms hydrogen bond interactions with residues H41, G143, S144, C145, H164 and E166 (Fig. 4d, e). The P1 pyridinyl group is positioned in the S1 region and forms hydrogen bond with His163. Compared to calpain inhibitor XII, in **UAWJ246** the 3-(pyrrolidin-2-one)methyl substitution occupies the S1 subsite instead of (2-pyridinyl)methyl in calpain inhibitor XII, leading to additional stabilizing hydrogen bonds as described previously. In addition, the small cyclopropyl group fits in S1’ subsite, avoiding steric repulsions with S1’ subsite amino acids as seen in the MD simulation trajectory with calpain inhibitor XII. These changes resulted in a potency enhancement by 100-fold, i.e., from 0.45 μΜ for calpain inhibitor XII to 0.045 μΜ for **UAWJ246**. In **UAWJ248** the length of the peptide was increased by adding a leucine between P2 Leu and Cbz group and additional lipophilic contacts with P168 are observed (Fig. 4p, q, s, t) but the activity remained unchanged. Two drug-bound complexes are shown for **UAWJ246** (pose 1 and pose 2) (Fig. 4g-i, 4j-l) and for **UAWJ248** (pose 1 and pose 2) (Fig. 4p-r, 4s-u), which correspond to different protomers. Minor differences are observed in the hydrogen bonding interactions between the two binding cavities in each protomer, reflecting the dynamic nature of the complexes. For example, a hydrogen bond with His41 was observed in **UAWJ246** pose 1 (Fig. 4g, h), while in **UAWJ246** pose 2 a hydrogen bond with Asn142 was observed (Fig. 4j, k). Similarly, hydrogen bonds with His41 and Thr190 were observed in **UAWJ248** pose 1 (Fig. 4p, q), and a hydrogen bond with Ser144 was observed in **UAWJ248** in pose 2 (Fig. 4s, t).

### Cellular antiviral activity and cytotoxicity of GC-376 analogues

To profile the antiviral activity of the **GC-376** analogues **UAWJ246**, **247**, and **248**, we first tested their cellular cytotoxicity against multiple cell lines. All three compounds were not toxic to these cell lines with CC_50_ values greater than 100 μM in most cases (Supplementary information Table S2). As such, we set the highest drug concentration as 30 μM or 100 μM in the antiviral assay. The antiviral activity of **GC-376** analogues was tested in both the immunofluorescence assay and plaque assay using the wild-type SARS-CoV-2 virus. **GC-376** was included as a positive control. In the immunofluorescence assay with the SARS-CoV-2 GFP reporter virus, **GC-376**, **UAWJ246**, **UAWJ247**, and **UAWJ248** inhibited the viral replication in a dose-response manner with EC_50_ values of 1.50 ± 0.42 μM, 15.13 ± 6.44 μM, 6.81 ± 0.65 μM, and 20.49 ± 3.71 μM, respectively (Fig. 5a-d, 5i). In the plaque assay, **GC-376**, **UAWJ246**, **UAWJ247**, and **UAWJ248** inhibited the viral replication with EC_50_ values of 0.48 ± 0.29 μM, 4.61 ± 3.60 μM, 2.06 ± 0.93 μM, and 11.1 ± 4.2 μM, respectively (Fig. 5e-h, 5j). Overall, all three **GC-376** analogues **UAWJ246**, **UAWJ247**, and **UAWJ248** had confirmed antiviral activity in cell culture. Comparing M^pro^ binding and anti-viral potency among the **GC-376** analogues, it appears that the aldehyde warhead may be more suitable for cell-based activities than the α-ketoamide. While the terminal groups of **UAWJ248** may enhance the enzymatic inhibition potency of **GC-376** by simultaneously occupying both S3 and S4 subpockets, the weaker cellular antiviral activity of **UAWJ248** might be due to decreased cellular permeability or increased metabolic degradation. In addition, as shown by our previous study, the calpain inhibitors demonstrated better antiviral activity than **GC-376** despite showing weaker binding affinity against M^pro^ in vitro,^15^ consistent with our hypothesis of synergistic inhibition of cathepsin L.

**Fig. 5.**
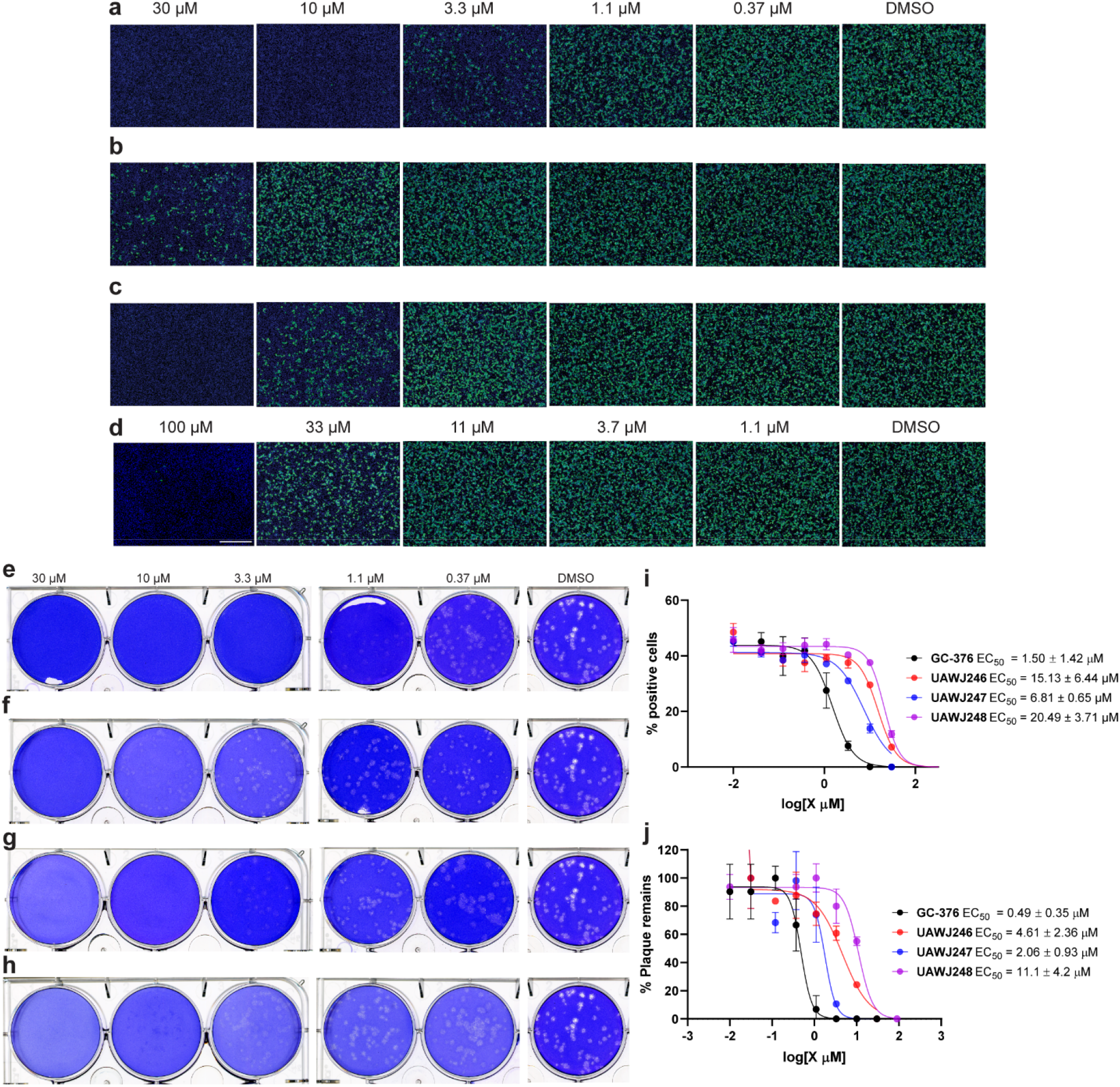
Antiviral activity of GC-376 analogues. **a-d** Antiviral activity of **GC-376** analogues against SARS-CoV-2 in the immunofluorescence assay. **a GC-376**; **b UAWJ246**; **c UAWJ247**; **d UAWJ248**. Vero E6 cells in a 96-well plate were infected with SARS-CoV-2 (USA-WA1/2020 isolate) at MOI of 0.05 in the presence of the indicated concentrations of the tested compounds. At 48 hpi, the cells were fixed and stained with a rabbit monoclonal antibody against the SARS-CoV-2 NP and a secondary antibody conjugated with Alexa 488 (Green). The nuclei were counterstained with Hoechst dye (Blue). For each well, fluorescence images of approximately 10K cells were acquired and shown. The images are representatives of two repeats. **e-h** Antiviral activity of **GC-376** analogues against SARS-CoV-2 in the plaque assay. **e GC-376**; **f UAWJ246**; **g UAWJ247**; **h UAWJ248**. Vero E6 cells in 6-well plates were infected with approximately 40 PFU/well of SARS-CoV-2 (USA-WA1/2020 isolate). After 1 hour, the inoculum was removed, and the cells were overlaid with medium containing the indicated concentrations of the tested compounds and 1.2% Avicel RC-591. At 3 dpi, the overlay was removed, and the cells were stained with 0.2% crystal violet. The images are representatives of two repeats. Data fitting of the antiviral activity of **GC-376** analogues against SARS-CoV-2 in the immunofluorescence assay (**i**) and the plaque assay (**j**).

## CONCLUSION

The ongoing COVID-19 pandemic needs an immediate intervention. If the previous SARS-CoV and MERS-CoV outbreaks are not severe enough to attract the attention from the scientific community, the current COVID-19 outbreak is a timely reminder of the threat of coronavirus. Encouraging progress has been made in developing antivirals and vaccines against SARS-CoV-2, such as remdesivir. However, despite the proof-reading function of the SARS-CoV-2 RdRp, SARS-CoV-2 continues to mutate, which will inevitably lead to drug resistance. Drug resistance has been evolved in cell culture against remdesivir using a model coronavirus, the murine hepatitis virus (MHV),^39^ raising concerns for the monotherapy of remdesivir. As such, new drugs with distinct mechanisms of action are needed.

The coronavirus M^pro^ (3CL^pro^) has long been pursued as a promising antiviral drug target.^12^ The unique feature of M^pro^ is its stringent preference for the glutamine residue at the P1 position, while no known host protease has such preference. Accordingly, the majority of M^pro^ inhibitors are designed to contain a glutamine mimetic at the P1 position such as pyrrolidone or 2-piperidinone. One of the most advanced lead compounds in this class is **GC-376**, an investigational veterinary drug that is currently being developed to treat feline infectious peritonitis. **GC-376** has optimal *in vivo* pharmacokinetic properties and *in vivo* antiviral efficacy in FIPD infection cat model.^40,41^ Our earlier study, coupled with an independent study from Vuong *et al*, showed that **GC-376** can similarly inhibit the enzymatic activity of SARS-CoV-2 M^pro^ and the viral replication of SARS-CoV-2 in cell culture.^37^ While this result is expected, our study also identified three additional non-conventional hits, Boceprevir, calpain inhibitors II and XII.^15^ These three compounds differ from known M^pro^ inhibitors in that they contain hydrophobic substitutions at the P1 site, challenging the notion that hydrophilic glutamine mimetics are required for potent inhibition. Intrigued by this finding, we pursued to solve the X-ray crystal structures of SARS-CoV-2 M^pro^ with Boceprevir and calpain inhibitors II and XII. During this process, the X-ray crystal structures of SARS-CoV-2 M^pro^ in complex with Boceprevir were released in PDB (PDB ID: 7BRP and 6WNP), and we therefore shifted our focus to calpain inhibitors II and XII. The binding pose of calpain inhibitor II in the active site of M^pro^ is consistent with other peptidomimetic inhibitors, where the methionine and leucine side chains occupy the S1 and S2 pockets, respectively. The methionine sulfur atom forms a hydrogen bond with the side chain imidazole of His163. In contrast, calpain inhibitor XII binds to the M^pro^ active site in an inverted conformation, projecting the pyridine instead of the norvaline residue in the S1 pocket. Again, the nitrogen from the pyridine forms a hydrogen bond with the side chain imidazole of His163. Collectively, these two structures suggest that His163 residue at the S1 pocket represents a binding hot spot for M^pro^ inhibitors, and that the pyridine ring is a suitable side chain to engineer potent interactions with the S1 subsite. The substrate plasticity of the P1 site suggests that it is feasible to design dual inhibitors targeting both the viral M^pro^ and other important proteases, such as cathepsin L. Cathepsin L has been identified as a critical host-protease for the SARS-CoV-2 cell entry. It plays an essential role in mediating the activation of the viral spike protein, thereby triggering membrane fusion and viral RNA release.^20,22^ Although the antiviral potency of cathepsin L inhibitors has been demonstrated against coronaviruses including SARS-CoV-2 in several studies, the *in vivo* antiviral efficacy of cathepsin L is not known. One potential concern is that the coronavirus spike protein can also be activated by other host proteases including trypsin, calpain, and TMPRSS2. Therefore, it is not clear whether inhibition of cathepsin L alone will be sufficient to completely stop viral replication *in vivo*. In this regard, a dual inhibitor design strategy that targets both the viral M^pro^ and host cathepsin L has certain advantages. First, compared to mono-specific M^pro^ inhibitors, dual inhibitors might have a higher genetic barrier to drug resistance as they also target the host cathepsin L. Second, compared to mono-specific cathepsin L inhibitors, dual inhibitors can lead to complete inhibition of viral replication as it targets the essential viral protein M^pro^.

Comparing X-ray crystal structures of SARS-CoV-2 M^pro^ in complex with **UAWJ246**, **UAWJ247**, and **UAWJ248** with the complex structure of **GC-376**, we can conclude that 1) the P1’ substitution does not contribute significantly to the potency of drug binding; 2) the P2 position prefers hydrophobic substitutions such as the leucine and phenylalanine side chains. Previous studies also demonstrate the cyclopropyl and cyclohexyl groups can be similarly accommodated.^17,18^ 3) the P3 and P4 positions prefer to be hydrophobic substitutions. However, these two positions, especially P3, may not be as important as the P1 and P2 substitutions, but favorable interactions with S3 and S4 subpockets can still contribute to the potency of inhibitor binding; 4) while the unique conformation of calpain inhibitor XII demonstrates the versatility of the α-ketoamide adduct formation, the antiviral activities of the **GC-376** analogues indicate that aldehyde-based compounds may be better suited for cell-based activity.

In summary, the structure-activity relationship of M^pro^ inhibitors revealed by the X-ray crystal structures and enzymatic assays described herein can be used to guide lead optimization. P1 substitution and the reactive warhead contribute significantly to the drug binding potency, followed by P2 substitution. While P1’, P3 and P4 substitutions are less essential, they can be optimized for inhibition against other proteases important for SARS-CoV-2 entry/replication and to improve their pharmacokinetic properties.

## MATERIALS AND METHODS

### Cell lines and viruses

Human rhabdomyosarcoma (RD), MDCK, Vero, Huh-7, and HCT-8 cell lines were maintained in Dulbecco’s modified Eagle’s medium (DMEM) medium; Caco-2 and MRC-5 cell lines were maintained in Eagle’s Minimum Essential Medium (EMEM) medium. Both media were supplemented with 10% fetal bovine serum (FBS) and 1% penicillin-streptomycin antibiotics. Cells were kept at 37°C in a 5% CO_2_ atmosphere. VERO E6 cells (ATCC, CRL-1586) were cultured in Dulbecco’s modified Eagle’s medium (DMEM), supplemented with 5% heat inactivated FBS in a 37°C incubator with 5% CO_2_. SARS-CoV-2, isolate USA-WA1/2020 (NR-52281), was obtained through BEI Resources and propagated once on VERO E6 cells before it was used for this study. Studies involving the SARS-CoV-2 were performed at the UTHSCSA biosafety level-3 laboratory by personnel wearing powered air purifying respirators.

### Protein expression and purification

SARS CoV-2 main protease (M^pro^ or 3CL) gene from strain BetaCoV/Wuhan/WIV04/2019 was ordered from GenScript (Piscataway, NJ) in the pET29a(+) vector with E. coli codon optimization. The expression and purification of His-tagged SARS CoV-2 M^pro^ (M^pro^-His) was described as previously.^15^

For HM-M^pro^ expression and purification, the SARS-CoV-2 Mpro gene from strain BetaCoV/Wuhan/WIV04/2019 GenScript (Piscataway, NJ, USA) was inserted into pETGSTSUMO vecror.The plasmid was transformed into Rosetta™(DE3) pLysS Competent Cells (Novagen). A single colony was picked for overnight growth to inoculate 50 mL of LB broth with 50□μg/mL kanamycin and 35 μg/ mL chloramphenicol. 10 mL of the overnight culture was used to inoculate 1 L of LB broth with 50 μg/mL kanamycin and 35 μg/ mL chloramphenicol. The 1L culture was grown at 250 RPM, 37 °C until OD 0.6~0.8. Expression was then induced with 0.5 mM IPTG at 250 RPM, 20 °C overnight. The culture was centrifuged at 5,000□g for 20 minutes and the resulting pellet was resuspended in 30□mL of the lysis buffer (20□mM Tris-HCl pH 8.4, 300□mM NaCl, 10% glycerol v/v, and 20□mM imidazole). These cells were lysed by sonication on a 10□second sonication/15□second rest cycle for a total of 15□minutes at an amplitude of 6. The lysate was centrifuged at 40,000 × g for 45□minutes at 4□°C and the supernatant was filtered, then loaded onto a HiTrap His column. The column was washed with lysate buffer and the protein was then eluted by linear gradient of imidazole. The peak of the protein was pulled and concentrated. The protein was then diluted in ULP1 cleavage buffer (20 mM Tris pH 8.0, 100 mM NaCl and 10 % Glycerol). The protease ULP1 was added at 1:20 ratio with incubation at 20C for overnight. The sample was loaded to His column and the flow through containing the untagged Mpro was collected. The untagged Mpro was concentrated and loaded to Superdex 200/16 equilibrated with 20 mM Tris pH 8.0, 250 mM NaCl. The peak of the protein was pooled and concentrated. The purity of the protein was evaluated by SDS-PAGE.

The expression and purification of SARS CoV-2 M^pro^ with unmodified N- and C-termini (M^pro^). SARS CoV-2 M^pro^ gene was subcloned from pET29a(+) to pE-SUMO vector according to manufacturer’s protocol (LifeSensors Inc, Malvern PA). pE-SUMO plasmid with SARS CoV-2 Main protease gene (M^pro^) was transformed into BL21(DE3) cells with kanamycin selection. A single colony was picked to inoculate 10 ml LB media and was cultured 37 °C overnight. This 10 ml culture was added to 1 liter LB media and grown to around OD 600 of 0.8. This culture was cooled on ice for 15 min, then induced with 0.5 mM IPTG. Induced cultures were incubated at 18°C for an additional 24 h and then harvested, lysed same way as His-tagged M^pro^ protein.^15^ The supernatant was incubated with Ni-NTA resin for overnight at 4 °C on a rotator. The Ni-NTA resin was thoroughly washed with 30 mM imidazole in wash buffer (50 mM Tris [pH 7.0], 150 mM NaCl, 2 mM DTT), SUMO-tagged M^pro^ was eluted from Ni-NTA with 300 mM imidazole. Eluted SUMO-tagged M^pro^ was dialyzed against 100-fold volume dialysis buffer (50 mM Tris [pH 7.0], 150 mM NaCl, 2 mM DTT) in a 10,000-molecular-weight-cutoff dialysis tubing. After dialysis, SUMO-tagged M^pro^ was incubated with SUMO protease 1 at 4 °C for overnight, and SUMO tag was removed by application of another round of Ni-NTA resin. The purity of the protein was confirmed with SDS-page gel.

The expression and purification of SARS CoV-2 papain-like protease (PL^pro^). SARS CoV-2 papain-like protease (PL^pro^) gene (ORF 1ab 1564 to 1876) from strain BetaCoV/Wuhan/WIV04/2019 was ordered from GenScript (Piscataway, NJ) in the pET28b(+) vector with E. coli codon optimization. The expression and purification procedure is very similar as we express SUMO-M^pro^ protein as we described in above section except that lysis buffer and Ni-NTA wash and elution buffer in pH7.5 (50 mM Tris [pH 7.5], 150 mM NaCl, 2 mM DTT). Human liver Cathepsin L was purchased from EMD Millipore (Cat # 219402).

### Peptide synthesis

The SARS-CoV-2 M^pro^ FRET substrate Dabcyl-KTSAVLQ/SGFRKME(Edans) was synthesized as described before.^15^ The SARS-CoV-2 PL^pro^ FRET substrate Dabcyl-FTLRGG/APTKV(Edans) was synthesized by solid-phase synthesis through iterative cycles of coupling and deprotection using the previously optimized procedure.^42^ Specifically, chemmatrix rink-amide resin was used. Typical coupling condition was 5 equiv of amino acid, 5 equiv of HATU, and 10 equiv of DIEA in DMF for 5 minutes at 80 °C. For deprotection, 5% piperazine plus 0.1 M HOBt were used and the mixture was heated at 80°C for 5 minutes. The peptide was cleaved from the resin using 95% TFA, 2.5% Tris, 2.5% H_2_O and the crude peptide was precipitated from ether after removal of TFA. The final peptide was purified by preparative HPLC. The purify and identify of the peptide were confirmed by analytical HPLC (> 98% purity) and mass spectrometry. [M+2]^2+^ calculated 888.04, detected 888.80.

### Compound synthesis and characterization

Details for the synthesis procedure (Supplementary Scheme 1) and characterization for compounds **UAWJ257**, **UAWJ246**, **UAWJ247**, and **UAWJ248** can be found in the supplementary information.

### Native Mass Spectrometry

Prior to analysis, the protein was buffer exchanged into 0.2 M ammonium acetate (pH 6.8) and diluted to 10 μM. DTT was dissolved in water and prepared at a 400 mM stock. Each ligand was dissolved in ethanol and diluted to 10X stock concentrations. The final mixture was prepared by adding 4 μL protein, 0.5 μL DTT stock, and 0.5 μL ligand stock for final concentration of 4 mM DTT and 8 μM protein. Final ligand concentrations were 10 μM. The mixtures were then incubated for 10 minutes at room temperature prior to analysis. Each sample was mixed and analyzed in triplicate.

Native mass spectrometry (MS) was performed using a Q-Exactive HF quadrupole-Orbitrap mass spectrometer with the Ultra-High Mass Range research modifications (Thermo Fisher Scientific). Samples were ionized using nano-electrospray ionization in positive ion mode using 1.0 kV capillary voltage at a 150 °C capillary temperature. The samples were all analyzed with a 1,000–25,000 m/z range, the resolution set to 30,000, and a trapping gas pressure set to 3. Between 10 and 50 V of source fragmentation was applied to all samples to aid in desolvation. Data were deconvolved and analyzed with UniDec.^43^

### Enzymatic assays

The main protease (M^pro^) enzymatic assays were carried out exact as previously described in pH 6.5 reaction buffer containing 20 mM HEPES pH6.5, 120 mM NaCl, 0.4 mM EDTA, 20% glycerol and 4 mM DTT.^15^

The SARS-CoV-2 papain-like protease (PL^pro^) enzymatic assays were carried out as follows: the assay was assembled in 96-well plates with 100 μl of 200 nM PL^Pro^ protein in PL^Pro^ reaction buffer (50 mM HEPES, pH7.5, 0.01% triton-100 and 5 mM DTT). Then 1 μl testing compound at various concentrations was added to each well and incubated at 30 °C for 30 min. The enzymatic reaction was initiated by adding 1 μl of 1 mM FRET substrate (the final substrate concentration is 10 μM). The reaction was monitored in a Cytation 5 image reader with filters for excitation at 360/40 nm and emission at 460/40 nm at 30 °C for 1 hr. The initial velocity of the enzymatic reaction with and without testing compounds was calculated by linear regression for the first 15 min of the kinetic progress curve. The IC_50_ values were calculated by plotting the initial velocity against various concentrations of testing compounds with a dose response function in Prism 8 software.

The cathepsin L enzymatic assay was carried out as follows: human liver cathepsin L (EMD Millipore 219402) was activated by incubating at reaction buffer (20 mM sodium acetate, 1 mM EDTA and 5 mM DTT pH5.5) for 30 min at 30 °C. Upon activation, the assay was assembled in 96-well plates with 100 μl of 300 pm cathepsin L protein in cathepsin L reaction buffer. Then 1 μl testing compound at various concentrations was added to each well and incubated at 30 °C for 30 min. The enzymatic reaction was initiated by adding 1 μl of 100 μM FRET substrate Z-Phe-Arg-AMC (the final substrate concentration is about 1 μM). The reaction was monitored in a Cytation 5 image reader with filters for excitation at 360/40 nm and emission at 460/40 nm at 30 °C for 1 hr. The IC_50_ values were calculated as described in above section.

### Differential scanning fluorimetry (DSF)

The thermal shift binding assay (TSA) was carried out using a Thermal Fisher QuantStudio™ 5 Real-Time PCR System as described previously.^15^ Briefly, 3 μM SARS-CoV-2 M^pro^ protein in M^pro^ reaction buffer (20 mM HEPES, pH 6.5, 120 mM NaCl, 0.4 mM EDTA, 4 mM DTT and 20% glycerol) was incubated with testing compounds at 30 °C for 30 min. 1X SYPRO orange dye was added and fluorescence of the well was monitored under a temperature gradient range from 20 °C to 90 °C with 0.05 °C/s incremental step.

### Cytotoxicity measurement

Evaluation of the cytotoxicity of compounds were carried out using the neutral red uptake assay.^44^ Briefly, 80,000 cells/mL of the tested cell lines were dispensed into 96-well cell culture plates at 100 μL/well. Twenty-four hours later, the growth medium was removed and washed with 150 μL PBS buffer. 200 μL fresh serum-free medium containing serial diluted compounds was added to each well. After incubating for 5 days at 37 °C, the medium was removed and replaced with 100 μL DMEM medium containing 40 μg/mL neutral red and incubated for 2-4 h at 37 °C. The amount of neutral red taken up was determined by measuring the absorbance at 540 nm using a Multiskan FC Microplate Photometer (Fisher Scientific). The CC_50_ values were calculated from best-fit dose response curves with variable slope in Prism 8.

### Molecular dynamics simulations

MD simulations were carried out to the covalently bound calpain inhibitor II, calpain inhibitor XII, **UAWJ246**, **UAWJ247**, **UAWJ248** with SARS-CoV-2 M^pro^ corresponding to PDB ID 6XA4 (monomer), PDB ID 6XFN (monomer), PDB ID 6XBG (dimer), PDB ID 6XBH (monomer), PDB ID 6XBI (dimer) prepared as described previously.^45^ The most favored protonation states of ionizable residues (D, E, R, K and H) at pH 7 were assigned using Maestro.^46^ The protonation states of the histidines in the binding region were set to δ position in order to contribute to the stabilization of complexes. Crystal waters were kept. All hydrogens atoms of the protein complex complex were minimized with the OPLS2005 force field^47^ by means of Maestro/Macromodel (Schrodinger 2017-1) using a distance-dependent dielectric constant of 4.0. The molecular mechanics minimization was performed with a conjugate gradient (CG) method and a root mean square of the energy gradient (threshold) value of 0.005 kJ Å^−1^ mol^−1^ was used as the convergence criterion. Each complex was solvated using the TIP3P^48^ water model. Using the “System Builder” utility of Schrodinger Desmond v.11.1 each complex was embedded in an orthorhombic water box extending beyond the solute 15 Å in x,y-plane and z-direction. Na^+^ and Cl^−^ ions were placed in the water phase to neutralize the systems and to reach the experimental salt concentration of 0.150 M NaCl. Each complex consists of c.a. 305 amino acid residues and 4,653 atoms and ~18,600 water residues (55,700 water atoms) or c.a. ~60,350 atoms for the monomer proteins and ~29,700 water residues (89,200 water atoms) for the dimer proteins, i.e., 98,000 atoms.

The OPLS2005 force field^49,50^ was used to model all protein and ligand interactions and lipids. The particle mesh Ewald method (PME)^51,52^ was employed to calculate long-range electrostatic interactions with a grid spacing of 0.8 Å. Van der Waals and short range electrostatic interactions were smoothly truncated at 9.0 Å. The Nose-Hoover thermostat was utilized to maintain a constant temperature in all simulations, and the Martyna-Tobias-Klein barostat was used to control the pressure. Periodic boundary conditions were applied (68×95×97) Å^3^ for the monomers and (105×106×96) Å^3^ for the dimers. The equations of motion were integrated using the multistep RESPA integrator^53^ with an inner time step of 2 fs for bonded interactions and non-bonded interactions within a cutoff 9 Å. An outer time step of 6.0 fs was used for non-bonded interactions beyond the cut-off.

Each system was equilibrated in MD simulations with a default protocol for water-soluble proteins provided in Desmond, which consists of a series of restrained MD simulations designed to relax the system, while not deviating substantially from the initial coordinates.

The first simulation was a Brownian dynamics run for 100 ps at a temperature of 10 K in the NVT (constant number of particles, volume, and temperature) ensemble with solute heavy atoms restrained with a force constant of 50 kcal mol Å^−2^. The Langevin thermostat^54^ was applied in the NVT ensemble and a MD simulation for 12 ps with solute heavy atoms restrained with a force constant of 50 kcal mol Å^−2^. The velocities were randomized and MD simulation for 12 ps was performed in the NPT ensemble and a Berendsen barostat^55^ with solute heavy atoms equally restrained at 10 K and another one at 300 K. The velocities were again randomized and unrestrained MD simulation for 24 ps was performed in the NPT ensemble. The above-mentioned equilibration was followed by 100ns simulation without restrains. Two MD simulations for each system were performed, one in in workstation with GTX 970, and using the GPU implementation of the MD simulations code, and one in ARIS-supercomputer system with cpu-cores. The visualization of produced trajectories and structures was performed using the programs Chimera^56^ and VMD.^57^

### Immunofluorescence assay

Vero E6 cells in 96-well plates (Corning) were infected with SARS-CoV-2 (USA-WA1/2020 isolate) at MOI of 0.05 in DMEM supplemented with 1% FBS. Immediately before the viral inoculation, the tested compounds in a three-fold dilution concentration series were also added to the wells in triplicate. The infection proceeded for 48 h without the removal of the viruses or the compounds. The cells were then fixed with 4% paraformaldehyde, permeabilized with 0.1% Triton-100, blocked with DMEM containing 10% FBS, and stained with a rabbit monoclonal antibody against SARS-CoV-2 NP (GeneTex, GTX635679) and an Alexa Fluor 488-conjugated goat anti-mouse secondary antibody (ThermoFisher Scientific). Hoechst 33342 was added in the final step to counterstain the nuclei. Fluorescence images of approximately ten thousand cells were acquired per well with a 10x objective in a Cytation 5 (BioTek). The total number of cells, as indicated by the nuclei staining, and the fraction of the infected cells, as indicated by the NP staining, were quantified with the cellular analysis module of the Gen5 software (BioTek).

### Plaque assay

Vero E6 cells in 6-well plates (Corning) were infected with SARS-CoV-2 (USA-WA1/2020 isolate) at approximately 40 PFU per well. After 1 hour of incubation at 37°C, the inoculum was removed and replaced with DMEM containing 1% FBS, 1.2% Avicel RC-591 (Dupont) and the tested compounds at different concentrations, in duplicate. After 3 days of infection, the overlay was removed, and the cells were fixed with 4% paraformaldehyde and stained with 0.2% crystal violet.

### M^pro^ crystallization and structure determination

SARS-CoV-2 M^pro^ was diluted to 5 mg/mL and incubated with 1.5 mM of inhibitor at 4 °C overnight. Samples were centrifuged at 13,000 G for 1 minute to remove precipitate. Crystals were grown by mixing the protein-inhibitor sample with an equal volume of crystallization buffer (20 % PEG 3000, 0.2 M Na Citrate pH 5.6) in a vapor diffusion, hanging drop apparatus. A cryoprotectant solution of 35 % PEG 3000 and 30 % glycerol was added directly to the drop and soaked for 15 minutes. Crystals were then flash frozen in liquid nitrogen for X-ray diffraction.

X-ray diffraction data for the SARS-CoV-2 M^pro^ structures were collected on the SBC 19-ID beamline at the Advanced Photon Source (APS) in Argonne, IL, and processed with the HKL3000 software suite. The CCP4 versions of MOLREP was used for molecular replacement using a previously solved SARS-CoV-2 M^pro^ structure, PDB ID: 7BRR as a reference model for the dimeric P2_1_ M^pro^ with UAWJ246. PDB ID 6YB7 was used as the reference model for the C2 monomeric M^pro^ with calpain inhibitors II/XII and UAWJ247, and the P1 dimeric structure with UAWJ248. PDB ID 6WTT was used as the reference model for the P3_2_21 trimer with UAWJ246. Rigid and restrained refinements were performed using REFMAC and model building was performed with COOT. Protein structure figures were made using PyMOL (Schrödinger, LLC).

## DATA AVAILABILITY

The drug-bound complex structures for SARS-CoV-2 M^pro^ have been deposited in the Protein Data Bank with accession numbers of 6XA4 (SARS-CoV-2 HM^Pro^ + Calpain inhibitor II), 6XFN (SARS-CoV-2 HM^Pro^ + Calpain inhibitor XII), 6XBG (SARS-CoV-2 HM^Pro^ + **UAWJ246**), 6XBH (SARS-CoV-2 HM^Pro^ + **UAWJ247**), and 6XBI (SARS-CoV-2 HM^Pro^ + **UAWJ248**).

## ACKNOWLEDGEMENTS

This research was partially supported by the National Institutes of Health (NIH) (Grant AI147325) and the Arizona Biomedical Research Centre Young Investigator grant (ADHS18-198859) to J. W. J. A. T. and M. T. M. were funded by the National Institute of General Medical Sciences and National Institutes of Health (Grant R35 GM128624 to M. T. M.). We thank Michael Kemp for assistance with crystallization and X-ray diffraction data collection. We also thank the staff members of the Advanced Photon Source of Argonne National Laboratory, particularly those at the Structural Biology Center (SBC), with X-ray diffraction data collection. SBC-CAT is operated by UChicago Argonne, LLC, for the U.S. Department of Energy, Office of Biological and Environmental Research under contract DE-AC02-06CH11357. The SARS-CoV-2 experiments were supported by a COVID-19 pilot grant from UTHSCSA to Y.X. SARS-Related Coronavirus 2, Isolate USA-WA1/2020 (NR-52281) was deposited by the Centers for Disease Control and Prevention and obtained through BEI Resources, NIAID, NIH.

## AUTHOR CONTRIBUTIONS

J. W. and Y. C. conceived and designed the study; C. M. expressed the M^pro^ and PL^pro^; C.M. performed the IC_50_ determination, thermal shift-binding assay, and enzymatic kinetic studies; M. S. carried out M^pro^ crystallization and structure determination with the assistance of X. Z, and analyzed the data with Y. C.; P. L. performed the molecular dynamics simulations under the guidance of A. K.; A. G. and N. K. synthesized the compounds for co-crystallization; Y. H. performed the cytotoxicity assay; X. M. and P. D. performed the SARS-CoV-2 immunofluorescence assay and plaque assay under the guidance of Y. X. B. H. and B. T. performed the initial antiviral assay with SARS-CoV-2; J. A. T. performed the native mass spectrometry experiments with the guidance from M. T. M.; J. W. and Y. C. secured funding and supervised the study; J. W., Y.C., and M.S. wrote the manuscript with the input from others.

## ADDITIONAL INFORMATION

Supplementary information accompanies this paper at

## Competing interests

The authors declare no competing interests.

## Notes

### Competing Interest Statement

The authors have declared no competing interest.

