## Supplementary information for "Structure and inhibition of the SARS-CoV-2 main protease reveals strategy for developing dual inhibitors against M^pro^ and cathepsin L"

| **Data Collection** | **PDB ID 6XA4** | **PDB ID 6XFN** | **PDB ID 6XBG** | **PDB ID 6XBH** | **PDB ID 6XBI** |
| --- | --- | --- | --- | --- | --- |
| Inhibitor | **Calpain inhibitor II** | **Calpain inhibitor XII** | **UAWJ246** | **UAWJ247** | **UAWJ248** |
| Space Group | C 1 2 1 | C 1 2 1 | P 1 21 1 | C 1 2 1 | P 1 |
| Cell Dimension |  |  |  |  |  |
| a, b, c (Å) | 113.538, 54.204, 45.414 | 114.566, 54.317, 44.492 | 55.367,98.854, 59.215 | 113.19,54.512, 45.129 | 46.898 54.903 62.261 |
| α, β, γ (°) | 90.00, 100.71, 90.00 | 90.00, 100.85, 90.00 | 90.00, 108.23, 90.00 | 90.00, 100.02, 90.00 | 60.63, 79.77, 88.83 |
| Resolution (Å) | 50.00 – 1.65 | 50.00 – 1.70 | 50.00 – 1.45 | 50.00 – 1.60 | 50.00 – 1.70 |
|  | (1.68 – 1.65) | (1.73 – 1.70) | (1.48 – 1.45) | (1.63 – 1.60) | (1.73 – 1.70) |
| R_merge_ | 0.065 (0.633) | 0.065 (0.129) | 0.086 (1.063) | 0.068 (0.500) | 0.083 (0.510) |
| <I>/σ<I> | 26.7 (1.33) | 29.34 (1.60) | 32.04 (2.36) | 25.82 (2.22) | 23.99 (2.36) |
| Completeness (%) | 98.2 (94.1) | 98.9 (99.8) | 98.7 (98.8) | 95.1 (93.9) | 98.2 (94.1) |
| Redundancy | 2.1 (1.9) | 4.1 (4.2) | 3.9 (3.7) | 3.4 (3.1) | 2.1 (1.9) |
| **Refinement** |  |  |  |  |  |
| Resolution (Å) | 50 – 1.65 | 50.00 – 1.70 | 50.00 – 1.45 | 50 – 1.60 | 50.00 – 1.70 |
|  | (1.71 - 1.65) | (1.76 - 1.70) | (1.50 - 1.45) | (1.66 - 1.60) | (1.76 - 1.70) |
| No. reflections/free | 32194 / 1596 | 29333 / 1490 | 105000 / 10263 | 33925 / 1673 | 52443 / 2619 |
| R_work_/R_free_ | 0.206 / 0.245 | 0.205 / 0.232 | 0.189 / 0.213 | 0.181 / 0.224 | 0.181 / 0.219 |
| No. Atoms | 2548 | 2561 | 5343 | 2668 | 5283 |
| Protein | 2356 | 2348 | 4822 | 2400 | 4798 |
| Ligand/Ion | 33 | 40 | 95 | 45 | 101 |
| Water | 159 | 173 | 426 | 223 | 384 |
| B-Factors (Å^2^) |  |  |  |  |  |
| Protein | 33.14 | 34.28 | 24.06 | 24.50 | 28.33 |
| Ligand/Ion | 36.26 | 43.73 | 30.080 | 25.80 | 29.36 |
| Solvent | 38.59 | 39.09 | 31.81 | 34.14 | 34.35 |
| RMS Deviations |  |  |  |  |  |
| Bond Lengths (Å) | 0.014 | 0.014 | 0.015 | 0.014 | 0.014 |
| Bond Angles (°) | 1.91 | 1.95 | 1.97 | 1.91 | 1.92 |
| Ramachandran Favored (%) | 98.68 | 98.34 | 98.16 | 98.68 | 97.20 |
| Ramachandran Allowed (%) | 1.32 | 1.66 | 1.84 | 1.32 | 2.80 |
| Ramachandran Outliers (%) | 0.00 | 0.00 | 0.00 | 0.00 | 0.00 |
| Rotameric Outliers (%) | 1.15 | 1.15 | 1.3 | 1.12 | 1.69 |
| Clashscore | 2.56 | 3.85 | 4.34 | 3.75 | 3.75 |

**Supplementary Table 1. Table of Crystallization Statistics.**

**Supplementary information Table S2: Cytotoxicity of SARS-CoV-2 M^pro^ inhibitors on various cell lines.**

|  | **UAWJ246**  CC_50_ (µM) | **UAWJ247**  CC_50_ (µM) | **UAWJ248**  CC_50_ (µM) |
| --- | --- | --- | --- |
| Caco-2 | > 250 | 146.3 ± 9.1 | 139.9 ± 14.4 |
| HCT-8 | > 250 | > 250 | > 250 |
| Huh-7 | > 250 | 108.2 ± 10.4 | > 250 |
| MRC-5 | > 250 | 98.4 ± 7.6 | 109.7 ± 9.8 |
| RD | > 250 | 93.2 ± 5.7 | 85.3 ± 7.8 |
| MDCK | > 250 | 214.4 ± 7.2 | 188.1 ± 3.6 |
| Vero | > 250 | 184.0 ± 4.8 | > 250 |

^a^Cytotoxicity was evaluated by measuring CC_50_ values (50% cytotoxic concentration) with CPE assay described in the method section. CC_50_ = mean ± S.E. of 3 independent experiments.

**Supplementary Scheme 1. Synthesis routes of UAWJ257 (a) and UAWJ248 (b).**

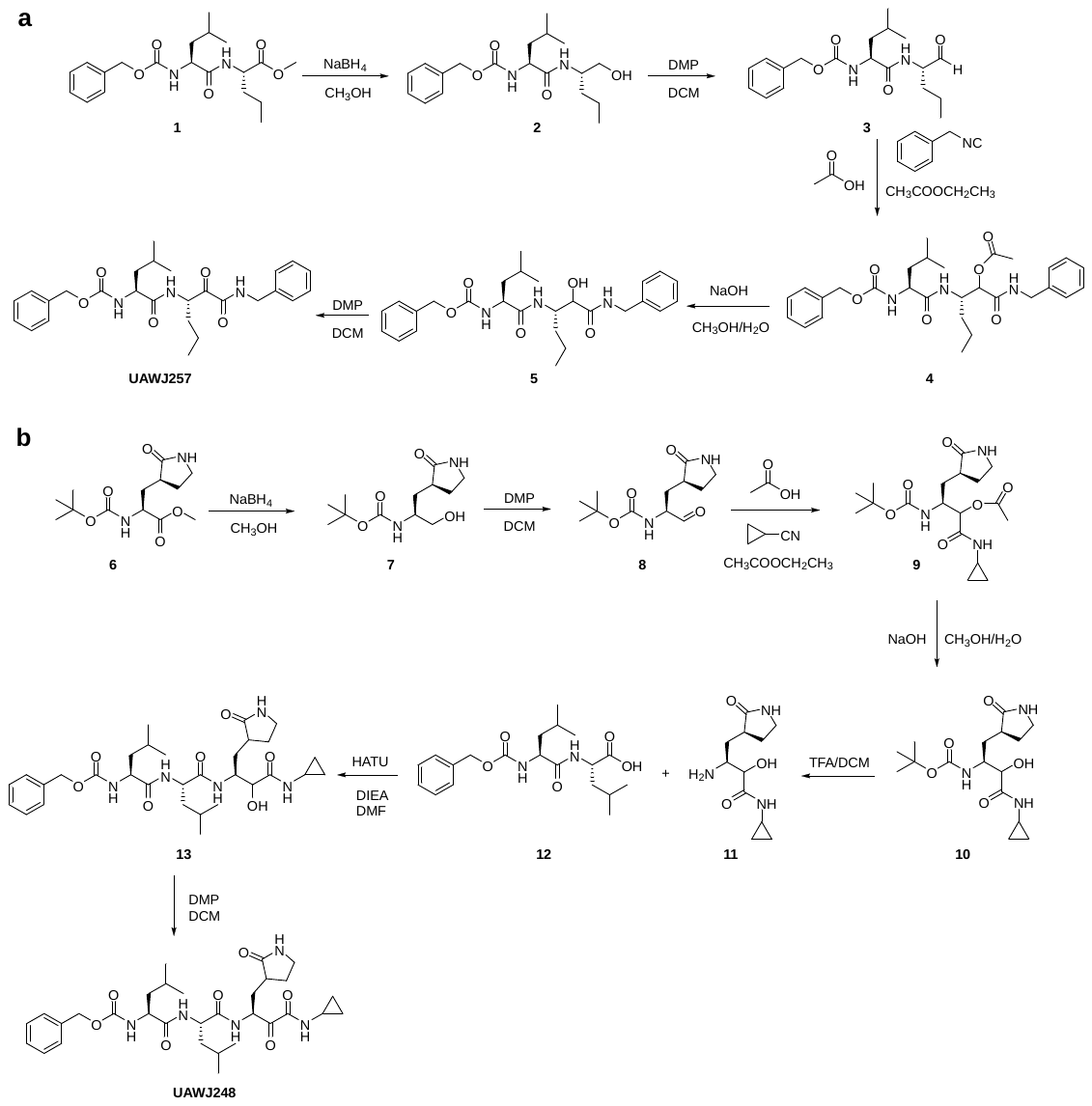

***General chemical methods.*** All chemicals were purchased from commercial vendors and used without further purification unless otherwise noted. ^1^H and ^13^C NMR spectra were recorded on a Bruker-400 or -500 NMR spectrometer. Chemical shifts are reported in parts per million referenced with respect to residual solvent (CD_3_OD) 3.31 ppm, (DMSO-d6) 2.50 ppm, and (CDCl_3_) 7.26 ppm or from internal standard tetramethylsilane (TMS) 0.00 ppm. The following abbreviations were used in reporting spectra: s, singlet; d, doublet; t, triplet; q, quartet; m, multiplet; dd, doublet of doublets; ddd, doublet of doublet of doublets. All reactions were carried out under N_2_ atmosphere, unless otherwise stated. HPLC-grade solvents were used for all reactions. Flash column chromatography was performed using silica gel (230–400 mesh, Merck). Low-resolution mass spectra were obtained using an ESI technique on a 3200 Q Trap LC/MS/MS system (Applied Biosystems). The purity was assessed by using Shimadzu LC-MS with Waters XTerra MS C-18 column (part #186000538), 50 × 2.1 mm, at a flow rate of 0.3 mL/min; λ = 250 and 220 nm; mobile phase A, 0.1% formic acid in H_2_O, and mobile phase B’, 0.1% formic in 60% isopropanol, 30% CH_3_CN and 9.9% H_2_O. All compounds submitted for testing in TEVC assay and plaque reduction assay were confirmed to be > 95.0% purity by LC-MS traces. All final products were characterized by proton and carbon NMR and MS.

***General procedure of the NaBH_4_ reduction.*** Compound **1** (10 mmol) was dissolved in 10 ml MeOH and the solution was cooled down to 0 °C. NaBH_4_ (10 eq.) was added slowly in portions. After stirring at ambient temperature for 1 hour, the solvent was partly removed *in vacuo* and 1 N HCl (aq.) was added. After 30 min stirring, EA was added. The organic phase was separated and dried with MgSO_4_. The solvent was removed *in vacuo* and purified with silica gel chromatography (EA/EtOH=5/1) to give compound **2** as a solid.

*benzyl N‐[(1S)‐1‐{[(2S)‐1‐hydroxypentan‐2‐yl]carbamoyl}‐3 methylbutyl]carbamate (****2****).* Yield: 92%. ^1^H NMR (400 MHz, CD_3_OD) δ 7.48 – 7.24 (m, 5H), 5.18– 5.07 (m, 2H), 4.19 (t, *J* = 7.6 Hz, 1H), 3.94 – 3.83 (m, 1H), 3.62 – 3.43 (m, 2H), 1.87 – 1.64 (m, *J* = 6.8 Hz, 1H), 1.64 – 1.51 (m, 3H), 1.52 – 1.22 (m, 3H), 1.08 – 0.83 (m, 9H). ^13^C NMR (101 MHz, CD_3_OD) δ 175.25, 158.38, 138.17, 129.42, 128.96, 128.78, 67.61, 65.02, 55.11, 52.30, 42.26, 34.12, 25.86, 23.44, 21.97, 20.16, 14.31.

***General procedure for the synthesis of aldehyde by Dess-Martin Periodinane (DMP) oxidation.*** Compound **2** was dissolved in anhydrous DCM and Dess-Martin periodinane (1.5 eq.) was added at 0 °C. The reaction mixture was stirred at ambient temperature for 1 hour then the solvent was removed *in vacuo*. 10% Na_2_S_2_O_3_ (aq.) was added and stirred for 30 min then EA was added. The organic phase was washed by 10% Na_2_S_2_O_3_ (aq.), saturated NaHCO_3_ (aq.), brine, dried by Na_2_SO_4_. The solvent was removed *in vacuo* and purified with silica gel chromatography (EA/EtOH=5/1) to give compound **3** as an oil. Due to the instability of the aldehyde, this intermediate was used immediately for the next step without further purification.

***General procedure for the Ugi three-component reaction.*** Aldehyde **3** (1mmol) was dissolved in ethyl acetate. Cyclopropyl isocyanide (1.2 eq.), HOAc (1.2 eq.) were added at 0 °C. The reaction mixture was stirred at ambient temperature for 12 hours until the starting martial disappeared. After removed the solvent *in vacuo*, the crude product was purified with silica gel chromatography (EA/EtOH=5/1) to give compound **4** as an oil. Compound **4** was used for the next step without further purification.

***General procedure for the hydrolysis of acetyl ester.*** Intermediate **4** (6 mmol) was dissolved in a mixture of H_2_O (18 mL) and MeOH (18 mL). NaOH (18 mmol) was added and the mixture was stirred 30 min. 1 N HCl (aq.) was added to adjust the pH to 6. The aqueous phase was extracted by chloroform 3 times, dried with NaSO_4_. The solvent was removed *in vacuo* to give intermediate **5** as a solid. Compound **5** was used for the next step reaction without further purification.

benzyl ((S)-1-(((S)-1-(benzylamino)-1,2-dioxohexan-3-yl)amino)-4-methyl-1-oxopentan-2-yl)carbamate (**UAWJ257**). **UAWJ257** was synthesized from the intermediate **5** by following the general procedure of DMP oxidation as described above. Yield: 64%. ^1^H NMR (400 MHz, DMSO-*d*_6_) δ 9.21 (t, *J* = 6.4 Hz, 1H), 8.24 (d, *J* = 6.7 Hz, 1H), 7.50 – 7.13 (m, 11H), 5.01 (s, 2H), 5.01 – 4.88 (m, 1H), 4.51 – 4.20 (m, 2H), 4.11 (q, *J* = 7.9 Hz, 1H), 1.89 – 1.57 (m, 2H), 1.53 – 1.17 (m, 5H), 0.94 – 0.76 (m, 9H). ^13^C NMR (101 MHz, DMSO-*d*_6_) δ 197.07, 172.63, 160.94, 155.88, 138.53, 137.10, 128.34, 127.78, 127.64, 127.30, 126.98, 65.35, 53.54, 52.63, 42.05, 40.80, 31.54, 24.12, 23.03, 21.56, 18.89, 13.54. C_27_H_35_N_3_O_5_, ESI-MS: m/z (M+H^+^): 482.6 (calculated), 482.6 (found).

Compounds **UAWJ246**, **UAWJ247** are known compounds and were synthesized according to reported procedures.^1,2^

*methyl (2S)‐2‐{[(tert‐butoxy)carbonyl]amino}‐3‐[(3S)‐2‐oxopyrrolidin‐3‐yl]propanoate (****6****).* Intermediate **6** was synthesized according to reported procedures.^3^

*tert‐butyl N‐[(2S)‐1‐hydroxy‐3‐[(3S)‐2‐oxopyrrolidin‐3‐yl]propan‐2‐yl]carbamate (****7****).* Intermediate **7** was synthesized from intermediate **6** by following the general procedure of NaBH_4_ reduction as described above. Yield: 80%. ^1^H NMR (400 MHz, CDCl3) δ 6.06 (s, 1H), 5.53 – 5.40 (m, 1H), 3.81 – 3.69 (m, 1H), 3.69 – 3.56 (m, 2H), 3.43 – 3.31 (m, 2H), 3.21 (s, 1H), 2.59 – 2.38 (m, 2H), 2.05 – 1.92 (m, 1H), 1.92 – 1.82 (m, 1H), 1.71 – 1.60 (m, 1H), 1.47 (s, 9H). ^13^C NMR (101 MHz, CDCl3) δ 180.78, 156.58, 79.48, 65.96, 51.12, 40.42, 38.01, 32.42, 28.45, 28.37.

*tert‐butyl N‐[(2S)‐1‐oxo‐3‐[(3S)‐2‐oxopyrrolidin‐3‐yl]propan‐2‐yl]carbamate (****8****).* Compound **8** was synthesized from intermediate **7** according to the general procedure of DMP oxidation as described above. Yield: 70%. ^1^H NMR (400 MHz, CDCl3) δ 9.52 (s, 1H), 6.83 (s, 1H), 6.12 (s, 1H), 4.21 – 4.07 (m, 1H), 4.07 – 3.96 (m, 1H), 3.39 – 3.24 (m, 2H), 2.51 – 2.29 (m, 2H), 1.92 – 1.65 (m, 2H), 1.39 (s, 9H).

*(2S)‐2‐{[(tert‐butoxy)carbonyl]amino}‐1‐(cyclopropylcarbamoyl)‐3‐[(3S)‐2‐oxopyrrolidin‐3‐yl]propyl acetate (****9****).* Compound **9** was synthesized from intermediate **8** according to the general procedure of the Ugi three-component reaction as described above. Yield: 72%. ^1^H NMR (400 MHz, CDCl_3_) δ 6.70 – 6.54 (m, 2H), 5.29 (d, *J* = 9.8 Hz, 1H), 5.19 – 5.09 (m, 1H), 4.22 – 4.01 (m, 1H), 3.39 – 3.25 (m, 2H), 2.73 – 2.62 (m, 1H), 2.42 (s, 2H), 2.15 (d, *J* = 6.9 Hz, 3H), 2.02 – 1.91 (m, 1H), 1.42 (s, 9H), 0.80 – 0.69 (m, 2H), 0.55 – 0.45 (m, 2H).

*tert‐butyl N‐[(2S)‐1‐(cyclopropylcarbamoyl)‐1‐hydroxy‐3‐[(3S)‐2‐oxopyrrolidin‐3‐yl]propan‐2‐yl]carbamate (****10****).* Compound **10** was synthesized from intermediate **9** according to the general procedure of acetyl ester hydrolysis as described above. Yield: 62%. ^1^H NMR (400 MHz, CDCl_3_) δ 7.06 – 6.93 (m, 1H), 5.51 – 5.38 (m, 1H), 4.22 – 4.01 (m, 2H), 3.45 – 3.29 (m, 2H), 2.80 – 2.67 (m, 1H), 2.60 – 2.48 (m, 1H), 2.47 – 2.35 (m, 1H), 2.13 – 1.98 (m, 1H), 1.96 – 1.83 (m, 1H), 1.83 – 1.66 (m, 1H), 1.44 (s, 9H), 0.84 – 0.74 (m, 2H), 0.59 – 0.51 (m, 2H). ^13^C NMR (101 MHz, CDCl_3_) δ 180.78, 173.62, 156.09, 79.64, 77.36, 73.03, 51.18, 40.51, 40.48, 37.77, 32.67, 28.31, 22.15, 6.42, 6.35.

*(3S)‐3‐amino‐N‐cyclopropyl‐2‐hydroxy‐4‐[(3S)‐2‐oxopyrrolidin‐3‐yl]butanamide (****11****).* Compound **10** (2 mmol) was treated by 4 M HCl in dioxane (2 ml) at 0 °C. After 1 hour at ambient temperature, the solvent was removed *in vacuo* to give compound **11** as a solid. The crude product was used directly without purification.

*benzyl N‐[(1S)‐1‐{[(1S)‐1‐{[(2S)‐1‐(cyclopropylcarbamoyl)‐1‐hydroxy‐3‐(2‐oxopyrrolidin‐3‐yl)propan‐2‐yl]carbamoyl}‐3‐methylbutyl]carbamoyl}‐3‐methylbutyl]carbamate (****13****).* Z-Leu-Leu-OH (1 mmol) and HATU (1.2 eq.) were dissolved in DMF (10 ml). After stirring for 30 min, compound **11** (1 eq.), DIEA (4 eq.) were added. The reaction mixture was stirred at ambient temperature for 20 hours. Then EA and water were added. The organic phase was washed by water, 1 M HCl (aq.), saturated NaHCO_3_ (aq.) and brine. The organic phase was dried by Na_2_SO_4_, and the solvent was removed *in vacuo* to give intermediate **13**. The crude product was used directly without purification.

*benzyl N‐[(1S)‐1‐{[(1S)‐1‐{[(2S)‐1‐(cyclopropylcarbamoyl)‐1‐oxo‐3‐(2‐oxopyrrolidin‐3‐yl)propan‐2‐yl]carbamoyl}‐3‐methylbutyl]carbamoyl}‐3‐methylbutyl]carbamate (****UAWJ248****).* Compound **UAWJ248** was synthesized from intermediate **13** according to the general procedure of DMP oxidation as described above. Yield: 55%. ^1^H NMR (400 MHz, DMSO-*d*_6_) δ 8.75 – 8.65 (m, 1H), 8.47 (d, *J* = 7.2 Hz, 1H), 7.94 – 7.80 (m, 1H), 7.65 (s, 1H), 7.47 – 7.24 (m, 6H), 5.10 – 4.91 (m, 3H), 4.40 – 4.26 (m, 1H), 4.04 (q, *J* = 8.1 Hz, 1H), 3.26 – 3.00 (m, 2H), 2.81 – 2.66 (m, 1H), 2.42 – 2.28 (m, 1H), 2.25 – 2.05 (m, 1H), 1.96 – 1.82 (m, 1H), 1.73 – 1.52 (m, 3H), 1.50 – 1.30 (m, 4H), 0.96 – 0.73 (m, 13H), 0.71 – 0.62 (m, 2H), 0.62 – 0.52 (m, 2H). ^13^C NMR (101 MHz, DMSO-*d*_6_) δ 196.32, 178.04, 172.29, 172.08, 162.14, 155.89, 137.10, 128.32, 127.74, 127.59, 65.31, 53.05, 52.08, 50.66, 40.99, 40.65, 39.42, 37.68, 31.05, 27.14, 24.18, 24.05, 23.04, 22.90, 22.48, 21.85, 21.49, 5.45, 5.42. C_31_H_45_N_5_O_7_, ESI-MS: m/z (M+H^+^): 600.7(calculated), 600.7 (found).

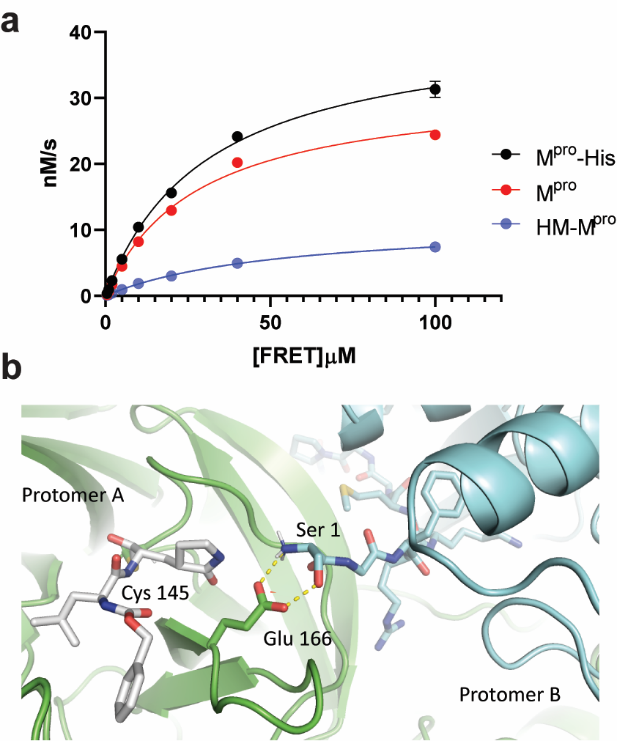

**Supplementary Figure 1.** Enzymatic activity and structure of SARS-CoV-2 M^pro^. **a** Michaelis-Menton plots of SARS CoV-2 His-tagged M^pro^ (M^pro^-His), Tag-free M^pro^ (M^pro^), and M^pro^ with extra His and Met residues at N-terminus (HM-M^pro^). The FRET substrate was added to 200 nM M^pro^-His, 200 nM M^pro^, or 1 µM HM-M^pro^ at different concentrations. The best fit V_max_ values are 41.4 ± 2.1 nM/s; 32.0 ± 2.4 nM/s, and 11.4 ± 0.9 nM/s for M^pro^-His, M^pro^, and HM-M^pro^ respectively; K_m_ values are 30.9 ± 3.8 µM, 27.8 ± 5.2 µM, and 53.1 ± 8.1 µM for M^pro^-His, M^pro^, and HM-M^pro^ respectively. The calculated *k*_cat_/K_m_ values for M^pro^-His, M^pro^ and HM-M^pro^ are 6,689 s^-1^M^-1^, 5,748 s^-1^M^-1^, and 214 s^-1^M^-1^ respectively. **b** Dimer structure of SARS-CoV-2 M^pro^ (PDB: 6WTT).

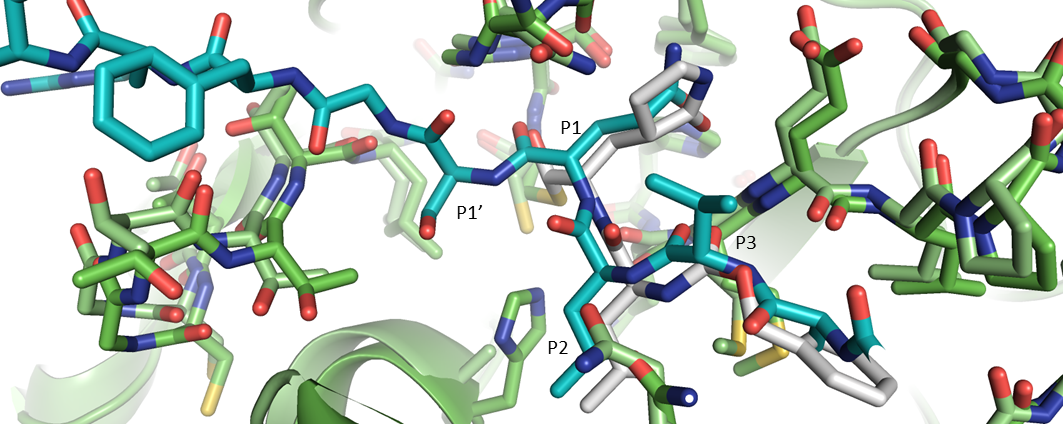

**Supplementary Figure 2**. Structure of catalytically impaired SARS-CoV M^pro^ H41A (light green) with N-terminal substrate (cyan) superimposed (2Q6G) with SARS-CoV-2 (dark green) bound to **GC-376** (6WTT, white).

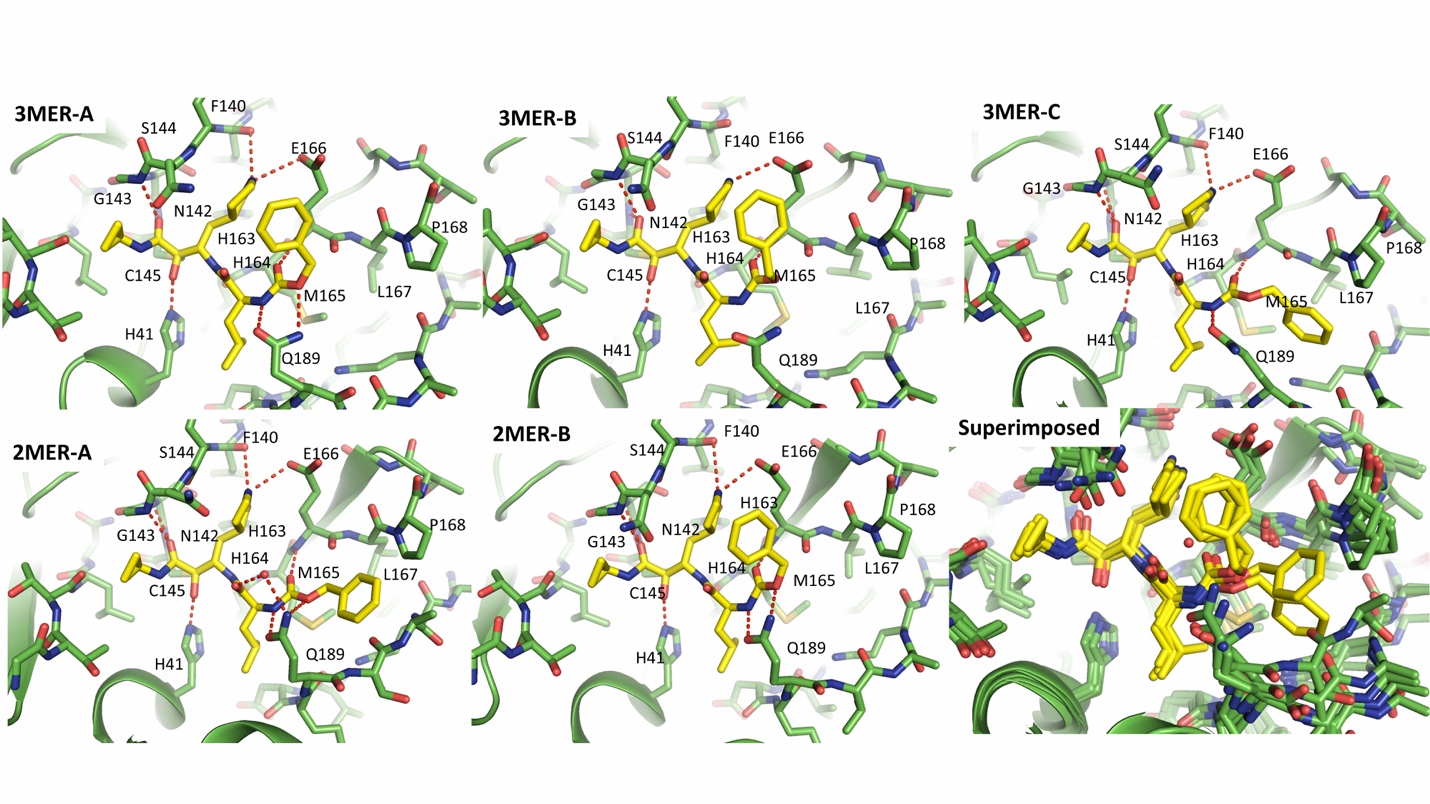
**Supplementary Figure 3.** Complex structure of **UAWJ246** with SARS-CoV-2 M^pro^-His in the P32_2_1 crystal form that contains three protomers per assymetric unit. Complex structure of **UAWJ246** with SARS-CoV-2 HM-M^pro^ in the dimer form was included for comparision.

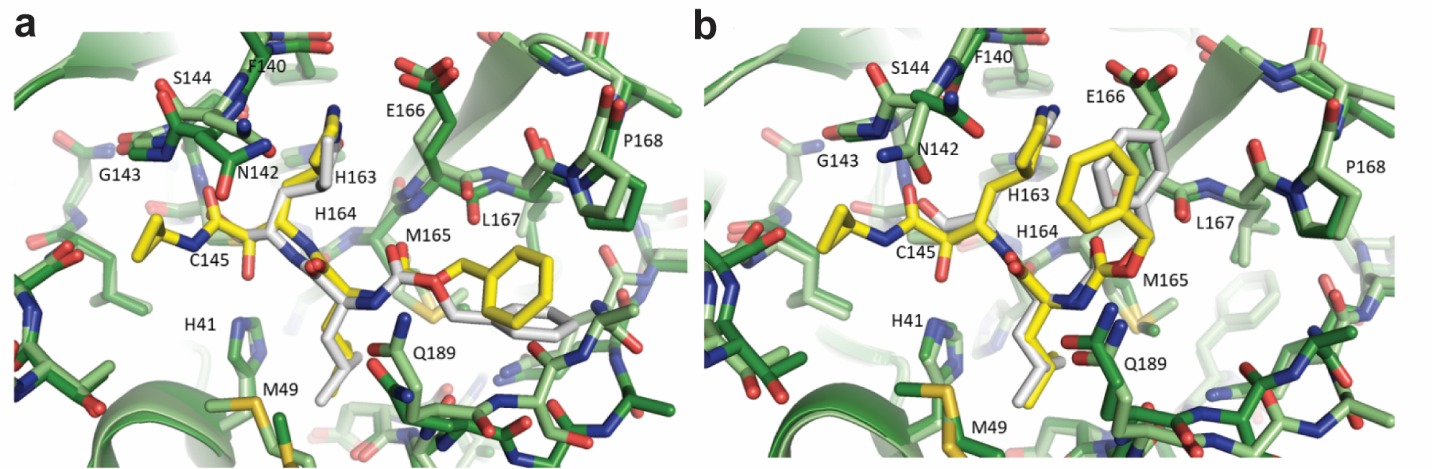

**Supplementary Figure 4.** Superimposition of SARS-CoV-2 HM-M^pro^ with **UAWJ246** (light green/yellow) and **GC-376** (dark green/grey). The conformation observed in the A protomer is consistent with the pose captured in PDB ID 6WTT (**a**).The conformation observed in the B protomer is consistent with the pose captured in PDB ID 7BRR (**b**).

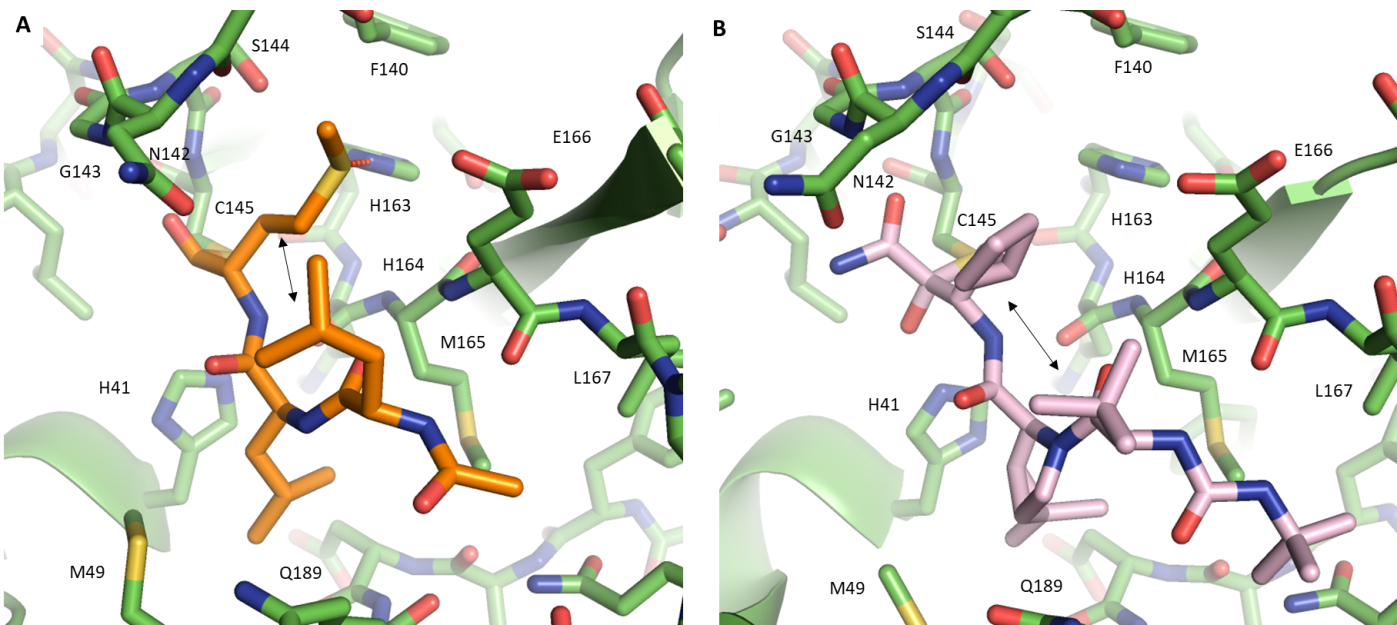

**Supplementary Figure 5.** Complex structure of SARS-CoV-2 HM-M^pro^ with **(A)** calpain inhibitor II and **(B)** Boceprivir (PDB ID 6WNP). In both structures a hydrophobic group is positioned in the P1 subsite. In the case of **(A)** calpain inhibitor II, a hydrogen bond is formed between the methionene sulfur and His 163. Intramolecular hydrophobic interactions between the S1 and S3 moieties are indicated by the arrow.
